# The role of Fragile X in the control of vertebrate spinal cord networks

**DOI:** 10.64898/2026.02.06.704392

**Authors:** Jonathan J. Milla-Cruz, Abel Mebrahtu, Laken Moller, Michelle A. Tran, Ning Cheng, Patrick J. Whelan

## Abstract

Fragile X syndrome (FXS) is the most common inherited cause of intellectual disability and the leading monogenic cause of autism, resulting from mutations in the *Fmr1* gene. While extensive research points to widespread circuit hyperexcitability across cortical and subcortical circuits, the contribution of the spinal cord circuits in the motor phenotypes associated with FXS remains largely unexplored. Given that *Fmr1* is expressed in both dorsal and ventral spinal cord, including motoneurons, the possibility exists that loss of its protein product, FMRP, disrupts locomotor circuitry. Here, we investigate whether *Fmr1* deletion alters the function of the spinal central pattern generator (CPG) networks and gait-related motor output. Using isolated neonatal spinal cord preparations from *Fmr1* knockout (*Fmr1* KO) mice, we assessed the ability of spinal circuits to generate coordinated fictive locomotor activity *in vitro*. In parallel, we quantified the gait parameters and motor performance in freely moving adult mice during unskilled and skill-demanding tasks. Our findings indicate that, despite the absence of FMRP in spinal neurons, neonatal *Fmr1* KO spinal cords generated robust and coordinated locomotor rhythms compared to controls. Consistently, adult *Fmr1* KO mice exhibited normal gait metrics under baseline conditions. However, these mice displayed hyperactivity and performance deficits during more challenging motor tasks demanding higher coordination. These findings suggest that the fundamental locomotor circuitry is preserved in FXS, likely through compensatory mechanisms. Consequently, motor impairments in FXS may arise primarily from supraspinal or integrative circuit dysfunction, rather than intrinsic deficits in spinal CPG function.

**Graphical Abstract:** 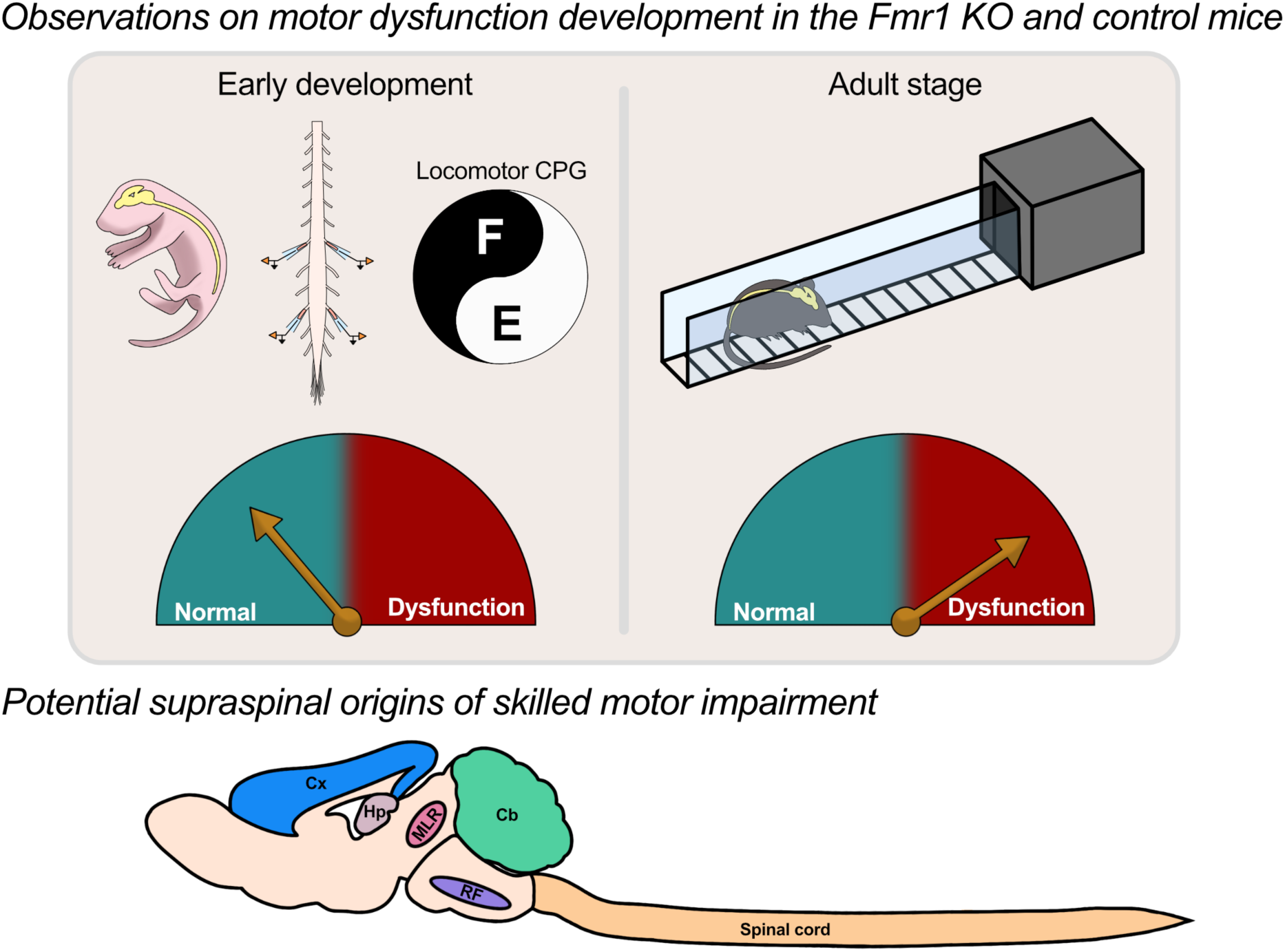

**Highlights:**

- Neonatal *Fmr1* KO spinal cords generated robust, coordinated locomotor rhythms similar to controls.
- Adult *Fmr1* KO mice exhibited normal gait metrics during baseline, unskilled locomotion.
- *Fmr1* KO mice displayed hyperactivity and performance deficits during skill-demanding motor tasks.
- FXS motor impairments may arise primarily from supraspinal or integrative circuit dysfunction
- Spinal cord circuitry appears to compensate for the fundamental loss of *Fmr1* function.

## Introduction

Fragile X syndrome (FXS) is caused by a mutation in the *Fmr1* gene on the X chromosome. As a result, males are more affected than females. Behaviorally, symptoms include anxiety, hyperactivity, and autism spectrum disorders ^1,2^. It is the most common cause of inherited intellectual disability and the leading cause of monogenic autism ^3^. Although *Fmr*1 is a single gene, its product, FMRP, binds a large set of activity-dependent neuronal mRNAs and controls their translation ^4^. Loss of FMRP dysregulates multiple synaptic and ion channels simultaneously ^5^. Because different brain and spinal regions rely on distinct FMRP-bound mRNA pools, *Fmr1* produces region-specific circuit abnormalities, making generalization difficult. That said, circuit-level abnormalities that have been most studied include hypersensitivity within the sensory system, linked to alterations at multiple levels of the neuraxis, including the spinal cord ^6–8^. In terms of hyperexcitability, somatosensory cortical neurons exhibit dysfunction of BK_Ca_ and I_h_, leading to increased firing rates ^9^. Hyperactivity is seen elsewhere, including in mouse hippocampus, cerebellum, and barrel cortex ^10–12^. Overall, this is due to a variety of channelopathies affecting plasticity, including long-term potentiation (LTP), upstates, and reduced tonic inhibition.

While the bulk of attention in terms of FXS has been on psychiatric dysfunction, there are also notable effects on motor function, which have both neural and muscular components. Infants and young children often exhibit low muscle tone, hyperextensible joints, and hip instability ^13,14^. In addition to hyperactivity, children with FXS show delays in acquiring skills such as independent walking. Older children often exhibit poor balance, coordination, and low endurance ^13^. These physical limitations exacerbate the social stigma associated with FXS, thereby increasing psychological issues. The *Fmr1* KO mouse model of FXS recapitulates many behavior-related characteristics, including hyperactivity and deficits in motor learning ^15–19^. Motor coordination dysfunction is observed in these mice during complex tasks, such as the balance beam ^20^. A limitation is that these studies were based on male mice, with less work on females. The expression of *Fmr1* in both the dorsal and ventral spinal cord, notably in motoneurons ^21^, is well documented in both mice and humans ^22,23^. Less is known about the function of *Fmr1* within the spinal cord. Although it is expressed in the dorsal horn and associated with sensory dysfunction, particularly in the nociceptive system ^22^, the ventral horn has received less attention. This observation raises questions about *Fmr1* and its role in modulating spinal cord circuits. The isolated spinal cord can produce basic locomotor patterns through central pattern generator circuits ^24,25^. Here, we examine whether the spinal central pattern generating network in *Fmr*1 KO mice is capable of producing coordinated locomotor output at birth using in vitro spinal cord preparations, and we also assess gait metrics in adult mice. Our data show that despite the absence of *Fmr1* from spinal cord neurons, there is no change in isolated locomotor output or adult gait patterns. The mice showed deficits in more complex motor tasks and exhibited hyperactivity, a characteristic of *Fmr1* mice. This suggests that, in terms of gait, the spinal cord may compensate for the loss of *Fmr*1.

## Results

### Analysis of the expression of FMRP in the brain and spinal cord of *Fmr1* KO mice

To determine the presence or absence of the FMRP in our *Fmr1* KO and FVB-wildtype (WT) mice, we performed an immunofluorescence assay using an antibody against the FMRP C-terminus. We examined expression in the spinal cord of adult mice (Figure 1). We observed a clear expression pattern in the spinal cord of FVB-WT mice in, but not in *Fmr1* KO mice.

**Figure 1:**
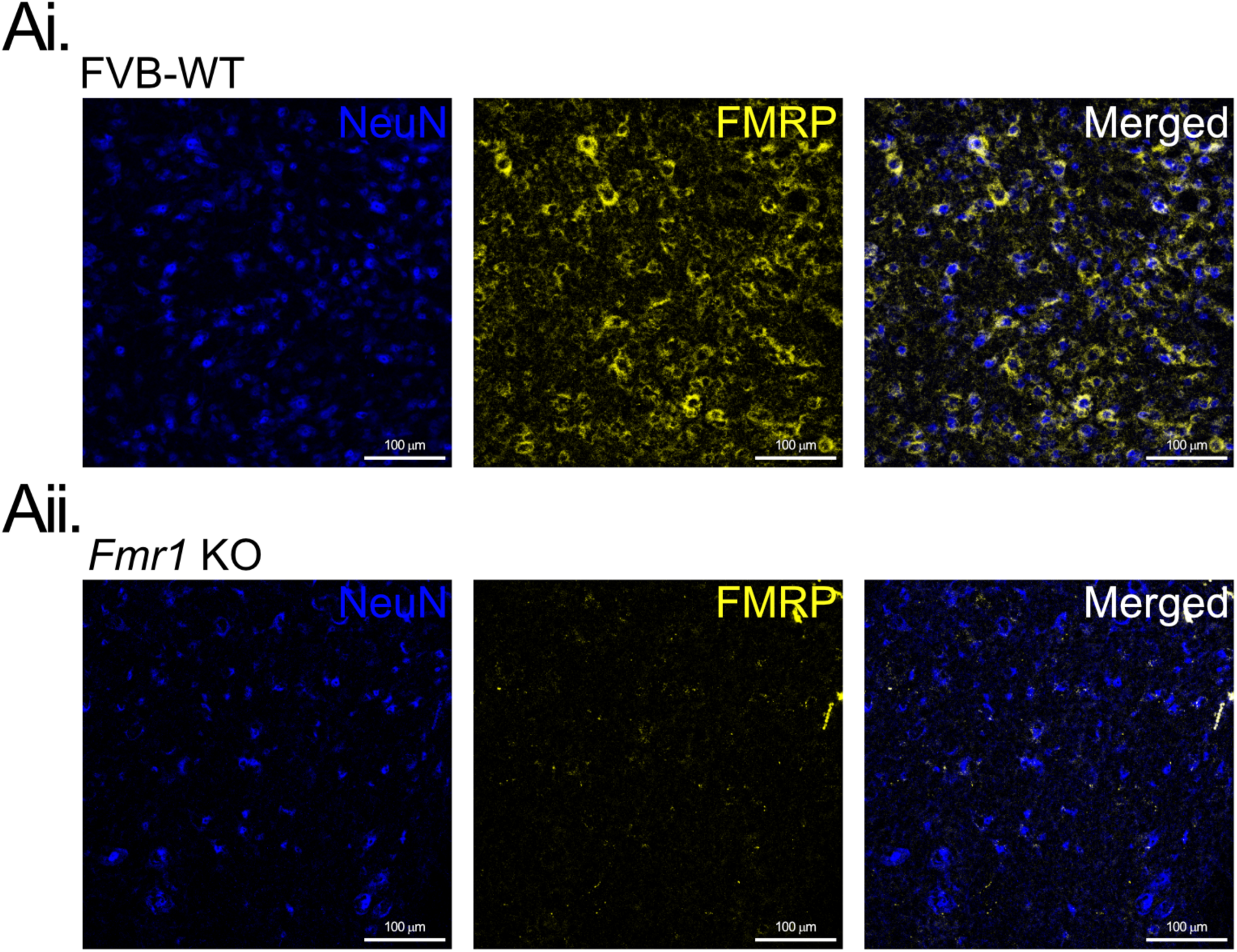
FMRP expression in the adult mouse brain and spinal cord. **A.** Representative transverse images from adult lumbar spinal cord sections expressing neuronal marker NeuN (blue) and FMRP (yellow) in FVB-WT (Ai) and *Fmr1* KO mice (Aii). Images depict the ventral horn area with a high density of FMRP-expressing neurons in the FVB-WT compared to *Fmr1* KO.

### Developmental milestones analysis in the *Fmr1* KO mice

Neurodevelopmental disorders such as FXS exhibit an early onset and progression during childhood; this makes the early developmental period a critical window not only for timely diagnosis, but also for the implementation of effective therapeutic strategies. Consequently, we studied the progression of FXS-related deficits in neonatal *Fmr1* KO mice.

To determine whether the *Fmr1* KO mice exhibit discernible delays in their neurobehavioral development compared to their control littermates, we examined their primitive reflexes at postnatal day 10 (P10) by using a battery of behavioral tests from previously established screens in mice ^26–28^. The rationale for selecting P10 to evaluate these reflexes is that the sensorimotor systems are mainly developed and mature by this time, allowing for a more accurate and comprehensive assessment of their functional integrity ^28^, Figure 2A). We first weighed each P10 mouse and then assessed the presence and, in some cases, the duration of eye opening, forelimb grasping, tactile reflexes (rooting, ear twitch), along with vestibular and postural reflexes (surface righting reflex, negative geotaxis, cliff aversion, and air righting reflex, Figure 2B).

**Figure 2:**
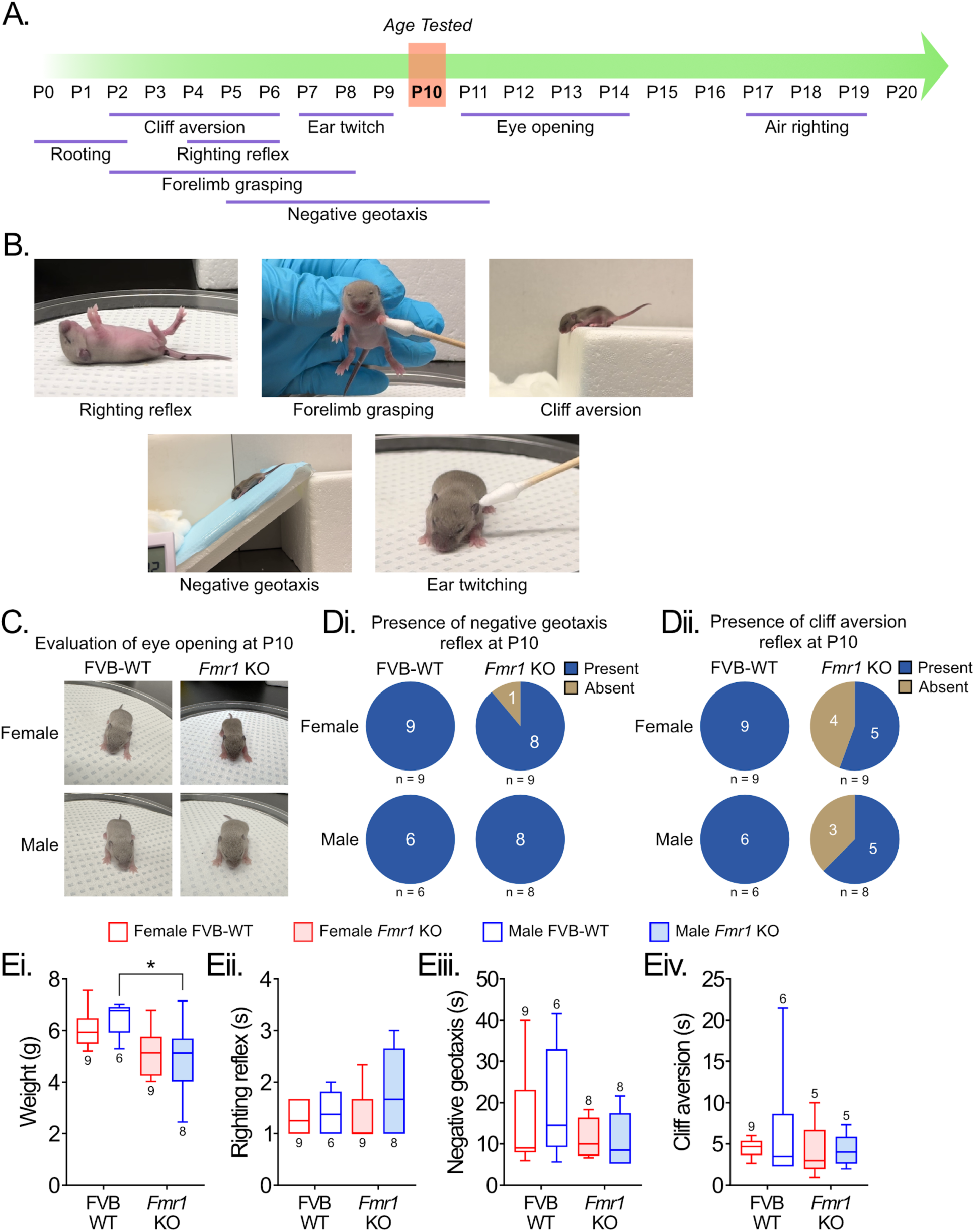
Examination of the developmental milestones in the *Fmr1* KO mice and FVB-WT mice at P10. **A.** Timeline indicating the appearance of sensorimotor reflexes in mice. Note that by P10 (tested age), most of the tested reflexes are known to be already present. **B.** Example photographs from pups engaged in the tests included in our battery of behavioral tests. **C.** Evaluation of the eye opening by P10 in both FVB-WT and *Fmr1* KO female and male pups. **D.** Evaluation of the presence or absence of the negative geotaxis reflex (Di) and cliff aversion reflex (Dii). Note that most of the pups showed the negative geotaxis by P10 for both FVB-WT and *Fmr1* KO, female and male mice, whereas only a fraction of the *Fmr1* KO female and male pups presented the cliff aversion reflex. **E.** Box and whisker plots display interquartile range (boxes), median (horizontal lines), maximum, and minimum values in data range (whiskers) for the analysis of the pup’s weight (Ei), vestibular and postural reflexes (Eii-Eiv) by P10. One-way ANOVA followed by Holm-Šídák multiple comparison test (*; *p* < 0.05).

Both groups of mice showed unopened eyes (Figure 2C). Forelimb grasping was present in both groups; negative geotaxis was present in all tested mice, except for 1 out of 8 female Fmr1 KO mice (Figure 2Di); and cliff aversion was present in all FVB-WT mice and in some *Fmr1* KO mice (Figure 2Dii). We observed that *Fmr1* KO pups weighed less than FVB-WT, with no difference in the female group (Figure 2Ei, Table 1, Supp. Table 1). However, the tactile reflexes (rooting, ear twitch) were present, whereas the air-righting reflexes were absent (data not shown).

**Table 1.**
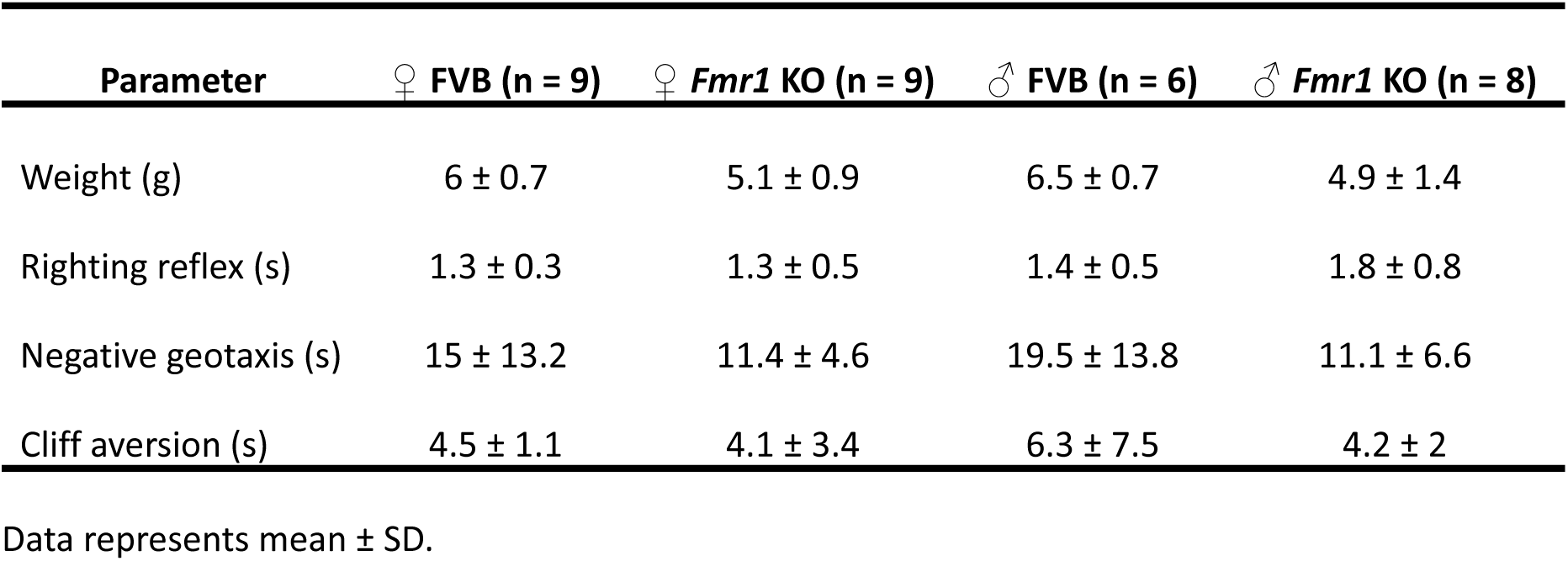
Behavioral analysis of the developmental milestones in the FVB-WT and *Fmr1* KO mice by P10.

Temporal analysis showed no differences in the righting reflex (Figure 2Eii, Supp. Table 1), negative geotaxis (Figure 2Eiii, Supp Table 1) nor cliff aversion reflex (Figure 2Eiv, Supp. Table 1), between *Fmr1* KO and FVB-WT in both female and male groups. These results suggest normal early sensorimotor development in *Fmr1* KO mice.

### Low-threshold sensory afferent transmission is not affected in the *Fmr1* KO mice during early development

Previous research has shown that ASD individuals with FXS can experience aberrant tactile discrimination and hypersensitivity to gentle touch during early development, leading to social interaction deficits and anxiety-like behaviors during adulthood ^29,30^. This somatosensory dysfunction is reported to result from a lack of GABA_A_ receptor-mediated presynaptic inhibition (PSI) of somatosensory neuron transmission in the spinal cord ^30^. For this reason, we explored the state of PSI on sensory afferent transmission in both mouse groups. We recorded the dorsal root potentials (DRPs) at the lumbar dorsal root L5 evoked by the stimulation of the tibial nerve (Tib) at different stimulus intensities (Figure 3A), as a way to measure the presynaptic inhibition on different modalities of afferent fibers ^31,32^. Since the amplitude and the area under the curve (AUC) of the DRPs depend on the seal’s stability from the suction electrode, we used the relative AUC and amplitude to analyze the differences between groups (see methods). We did not observe any significant difference in the peak amplitude, AUC or time to peak of the DRPs measured at any stimulus strength tested, between the FVB-WT and the *Fmr1* KO mice in female and male mice (Figure 3B, Table 2, Supp. Table 1). These data suggest that not only low-threshold afferent transmission (low stimulus intensities) but also high-threshold afferent transmission (nociceptive transmission) are not affected during the early development of the *Fmr1* KO mice.

**Figure 3:**
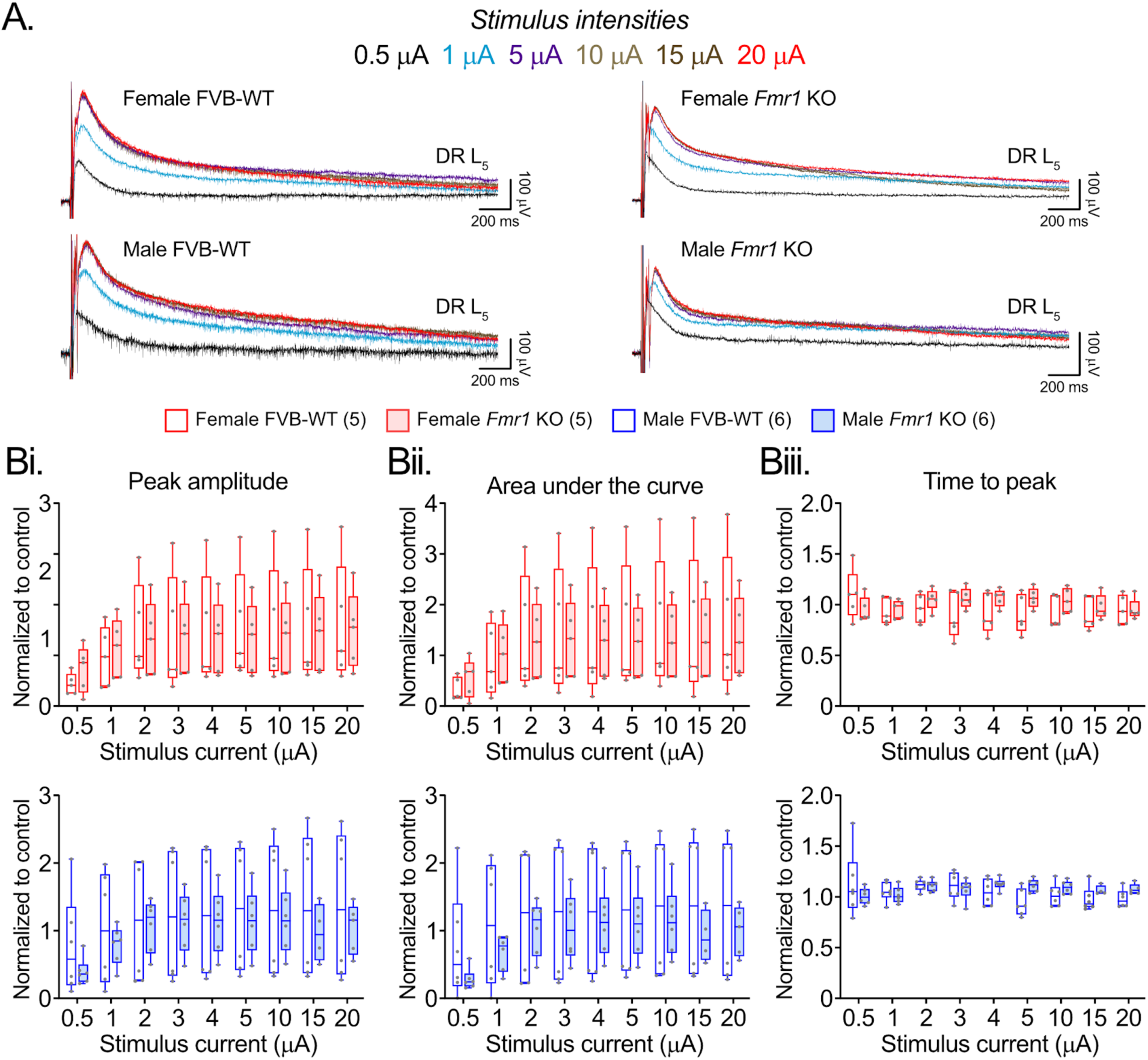
Sensory afferent transmission in the *Fmr1* KO mice is similar to FVB-WT. **A.** Representative traces of dorsal root potentials recorded at the lumbar L5 dorsal root entry zone, evoked by stimulation of the tibial nerve (Tib) at different stimulus intensities in the female and male FVB-WT and *Fmr1* KO mice. Six traces were averaged for each condition. **B.** Box and whisker plots display interquartile range (boxes), median (horizontal lines), maximum, and minimum values in data range (whiskers) for the analysis of the DRPs relative peak amplitude (Bi), area under the curve (300ms, Bii), and time to peak (Biii), evoked at different stimulus intensities. Two-way ANOVA followed by Šídák multiple comparison test (*; *p* < 0.05).

**Table 2.** Tib nerve evoked DRPs parameters in neonate FVB-WT and *Fmr1* KO mice.

| Stimulus (mA) | ♀ FVB (n = 5) | ♀ *Fmr1* KO (n = 5) | ♂ FVB (n = 6) | ♂ *Fmr1* KO (n = 6) |
| --- | --- | --- | --- | --- |
| *Peak amplitude* |  |  |  |  |
| 0.5 | 0.44 ± 0.21 | 0.73 ± 0.46 | 0.78 ± 0.74 | 0.40 ± 0.20 |
| 1 | 0.97 ± 0.60 | 1.14 ± 0.59 | 1.02 ± 0.83 | 0.79 ± 0.29 |
| 2 | 1.45 ± 0.96 | 1.32 ± 0.74 | 1.14 ± 0.92 | 1.08 ± 0.39 |
| 3 | 1.38 ± 1.17 | 1.36 ± 0.75 | 1.24 ± 0.99 | 1.21 ± 0.44 |
| 4 | 1.44 ± 1.14 | 1.35 ± 0.74 | 1.26 ± 0.98 | 1.13 ± 0.47 |
| 5 | 1.55 ± 1.10 | 1.33 ± 0.71 | 1.32 ± 0.95 | 1.14 ± 0.48 |
| 10 | 1.54 ± 1.18 | 1.37 ± 0.75 | 1.33 ± 1.01 | 1.16 ± 0.49 |
| 15 | 1.54 ± 1.20 | 1.43 ± 0.80 | 1.37 ± 1.07 | 0.97 ± 0.43 |
| 20 | 1.60 ± 1.20 | 1.49 ± 0.77 | 1.37 ± 1.08 | 1.03 ± 0.38 |
| *Area under curve* |  |  |  |  |
| 0.5 | 0.34 ± 0.23 | 0.55 ± 0.39 | 0.77 ± 0.82 | 0.29 ± 0.16 |
| 1 | 0.90 ± 0.72 | 1 ± 0.6 | 1.08 ± 0.93 | 0.68 ± 0.26 |
| 2 | 1.38 ± 1.17 | 1.29 ± 0.76 | 1.21 ± 1.01 | 1.04 ± 0.40 |
| 3 | 1.40 ± 1.30 | 1.32 ± 0.77 | 1.27 ± 1.06 | 1.05 ± 0.50 |
| 4 | 1.42 ± 1.34 | 1.29 ± 0.75 | 1.28 ± 1.02 | 1.12 ± 0.51 |
| 5 | 1.49 ± 1.29 | 1.27 ± 0.70 | 1.31 ± 0.99 | 1.12 ± 0.52 |
| 10 | 1.55 ± 1.34 | 1.29 ± 0.73 | 1.35 ± 1.04 | 1.14 ± 0.52 |
| 15 | 1.50 ± 1.41 | 1.34 ± 0.80 | 1.36 ± 1.06 | 0.92 ± 0.39 |
| 20 | 1.58 ± 1.40 | 1.37 ± 0.79 | 1.35 ± 1.06 | 1 ± 0.37 |
| *Time to peak* |  |  |  |  |
| 0.5 | 1.10 ± 0.25 | 0.95 ± 0.12 | 1.13 ± 0.33 | 1.01 ± 0.08 |
| 1 | 0.94 ± 0.13 | 0.96 ± 0.09 | 1.05 ± 0.10 | 1.02 ± 0.08 |
| 2 | 0.96 ± 0.14 | 1.05 ± 0.11 | 1.12 ± 0.06 | 1.11 ± 0.05 |
| 3 | 0.90 ± 0.23 | 1.06 ± 0.10 | 1.11 ± 0.14 | 1.07 ± 0.1 |
| 4 | 0.91 ± 0.20 | 1.07 ± 0.09 | 1.05 ± 0.13 | 1.13 ± 0.06 |
| 5 | 0.91 ± 0.19 | 1.07 ± 0.10 | 0.94 ± 0.13 | 1.11 ± 0.06 |
| 10 | 0.92 ± 0.16 | 1.04 ± 0.12 | 1.02 ± 0.11 | 1.09 ± 0.06 |
| 15 | 0.91 ± 0.16 | 0.98 ± 0.12 | 0.98 ± 0.12 | 1.07 ± 0.05 |
| 20 | 0.95 ± 0.15 | 0.95 ± 0.10 | 0.98 ± 0.09 | 1.08 ± 0.05 |

Data represents mean ± SD.

### Spontaneous activity showed differences between *Fmr1* KO and FVB-WT mice during early development

Spontaneous activity is important for the development of motor networks ^33^. Previous studies have shown that during early development, spinal cord networks express a distinctive motor output called spontaneous activity, characterized *in vitro* as sporadic episodes of uncoordinated rhythmic movements ^25,33^.

In this regard, we examined the spontaneous activity recorded from the lumbar ventral roots in neonatal *Fmr1* KO and FVB-WT mice. Subsequent analysis was performed using our previously developed SpontaneousClassification toolbox ^34^, which allows classification of single episodes based on their amplitude and rhythmicity into 5 different classes (Figure 4A). We observed that episode frequency and classification differed by sex and genotype (Figure 4Bi, Table 3, Supp. Table 1). Female *Fmr1* KO mice showed a greater number of large, non-rhythmic episodes (Class 2) compared to both female and male FVB-WT mice.

**Figure 4:**
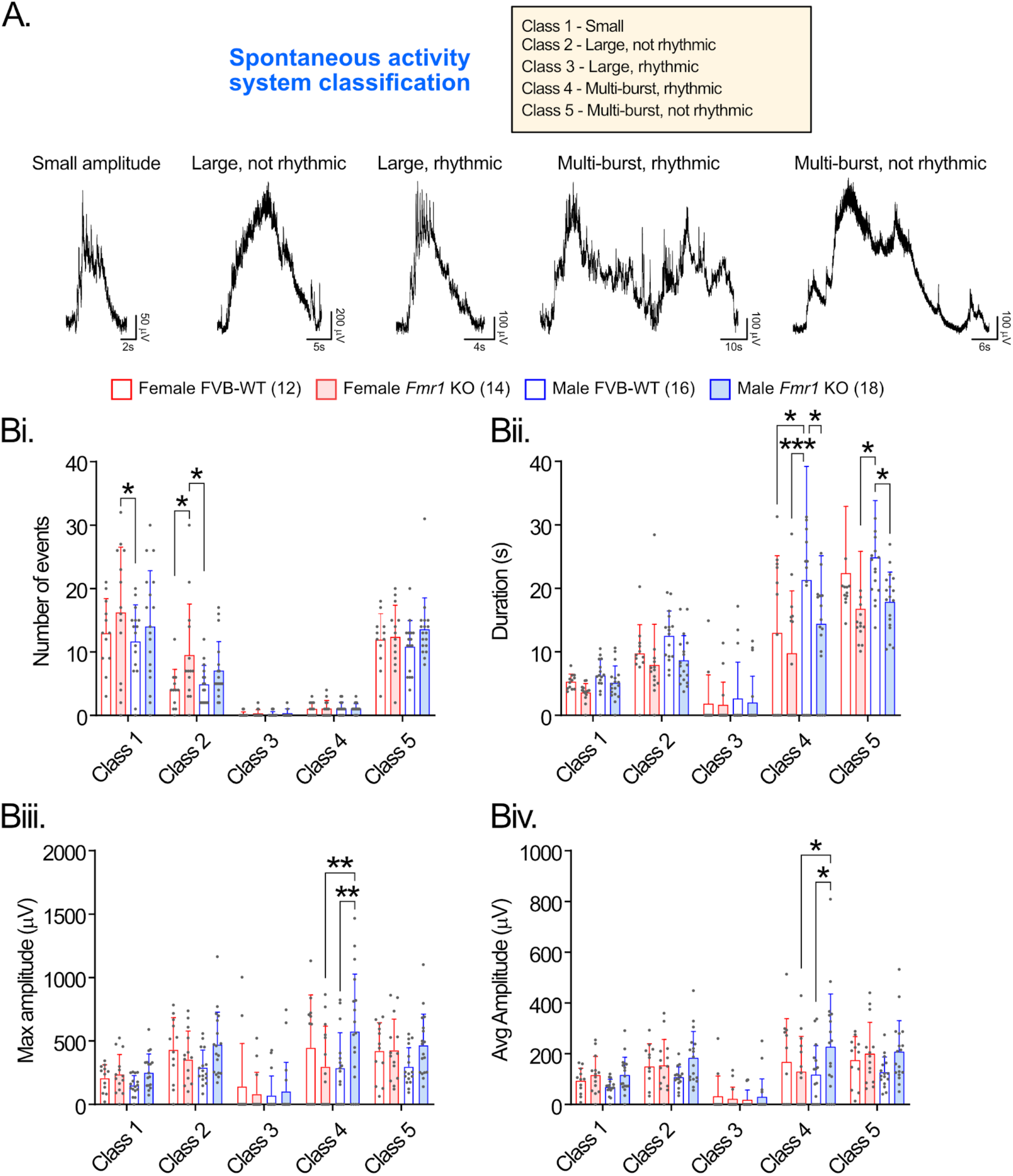
Analysis of the spontaneous activity in the *Fmr1* KO and FVB-WT control mice. **A.** Our previously developed classification system for spontaneous activity was used, sorting features into five different classes. These classes represent various characteristics, including amplitude, rhythmicity, and episodes of spontaneous activity. Representative traces show the main characteristics for each class. **B.** Bar plots for the analysis of the number of events (Bi), duration (Bii), max amplitude (Biii) and average amplitude (Biv) from classified episodes of spontaneous activity in the FVB-WT and *Fmr1* KO, female and male mice. Data are presented as mean ± SD with asterisks denoting significance (*: *p* < 0.05, **: *p* < 0.01, ***: *p* < 0.001) post hoc analyses following a two-way ANOVA.

**Table 3.**
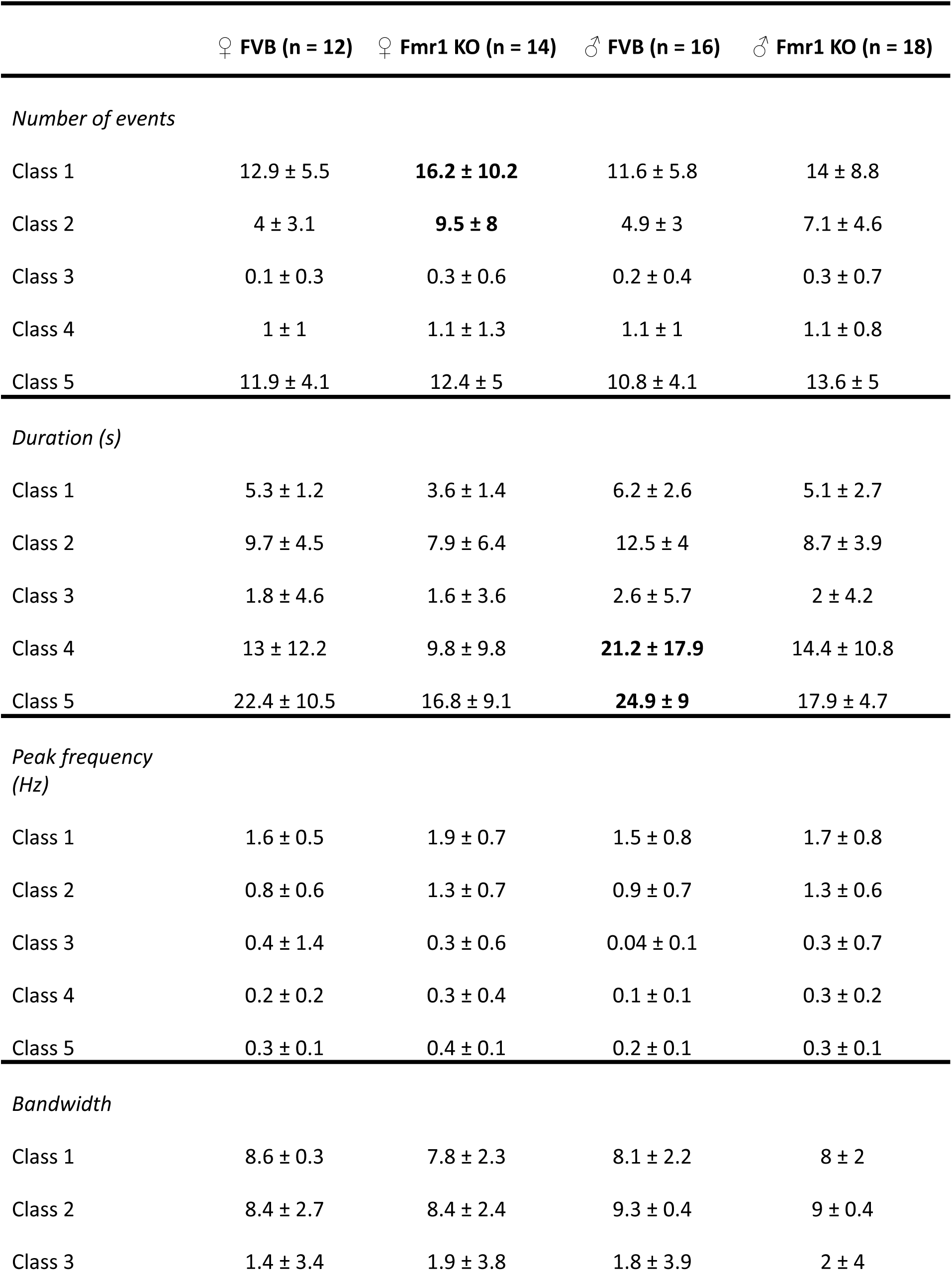
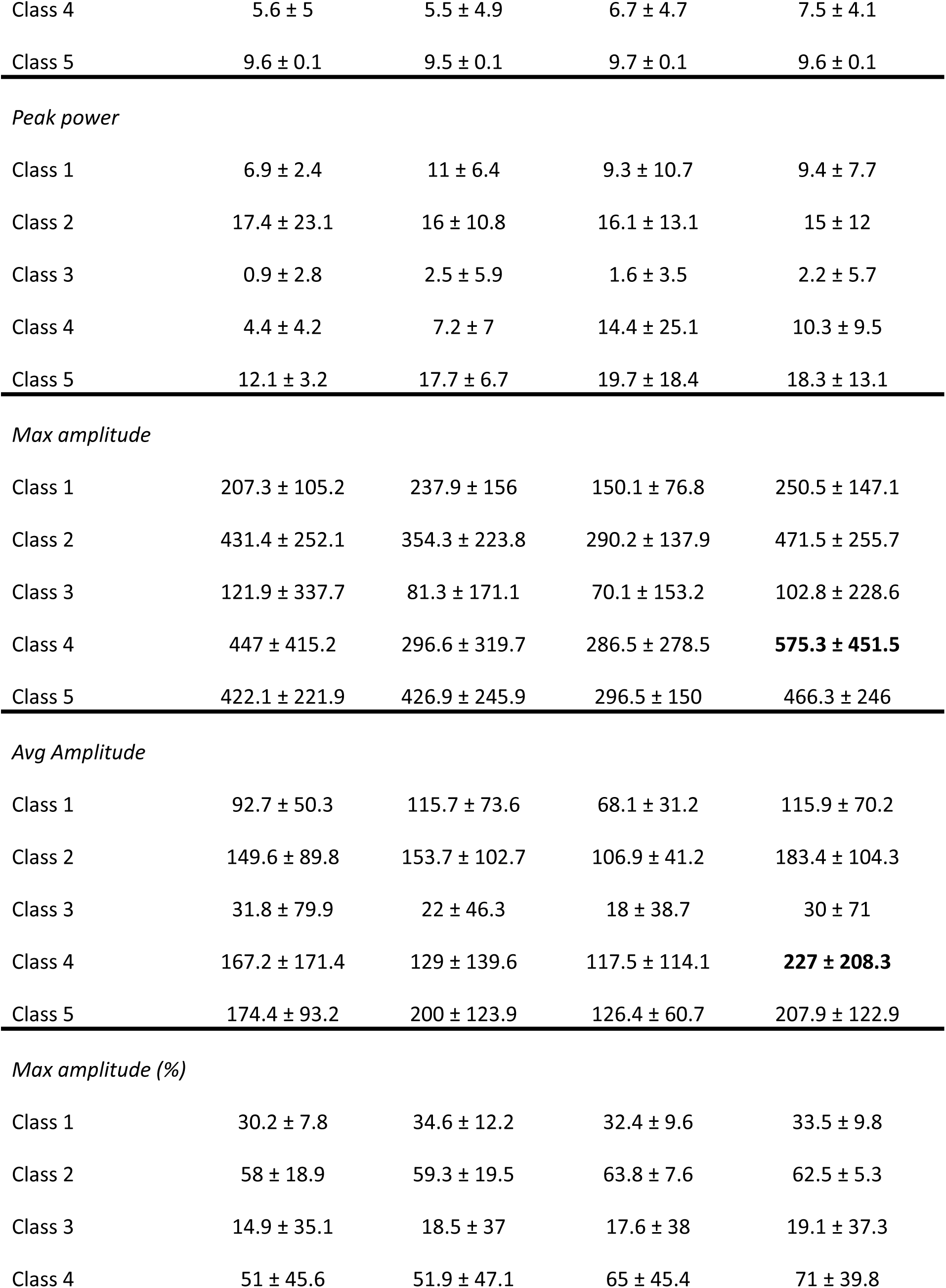
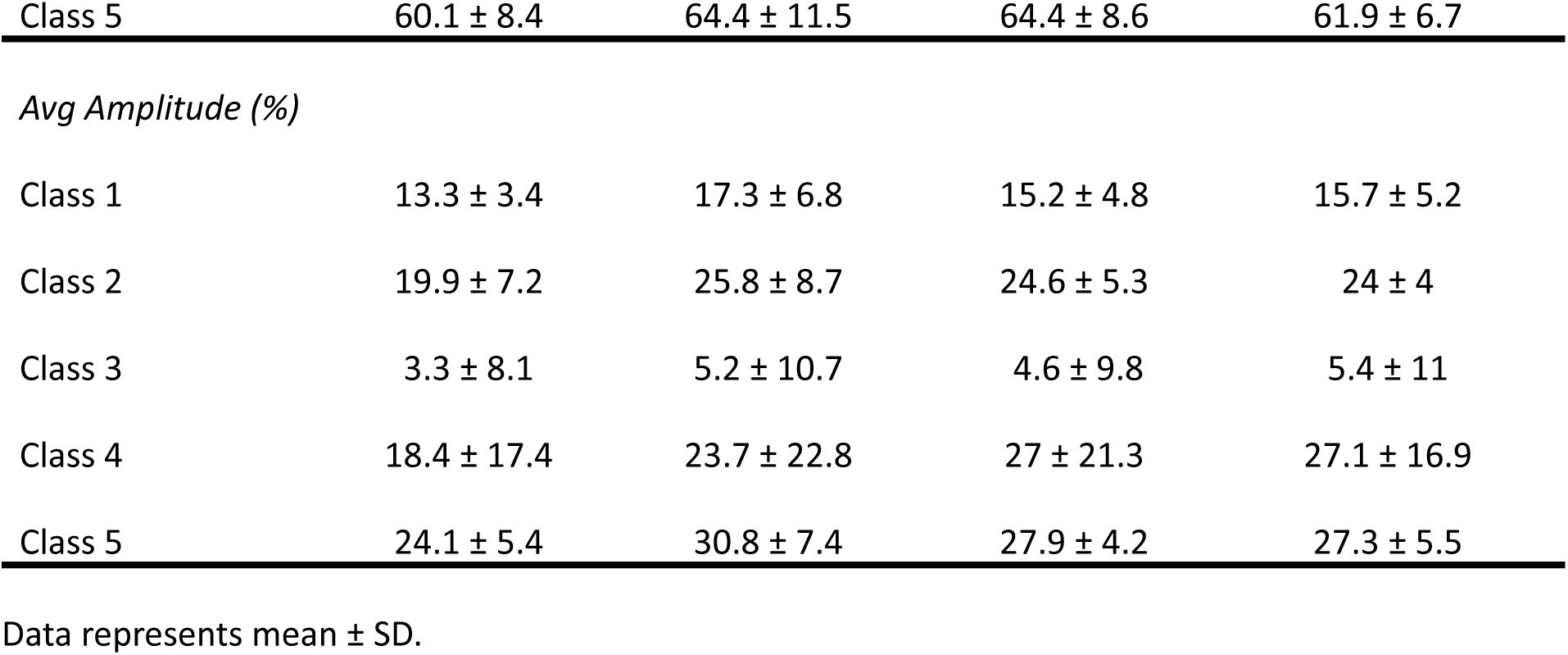
Spontaneous activity classification in FVB-WT and *Fmr1* KO mice.

|  | ♀ FVB (n = 12) | ♀ *Fmr1* KO (n = 14) | ♂ FVB (n = 16) | ♂ *Fmr1* KO (n = 18) |
| --- | --- | --- | --- | --- |
| *Number of events* |  |  |  |  |
| Class 1 | 12.9 ± 5.5 | **16.2 ± 10.2** | 11.6 ± 5.8 | 14 ± 8.8 |
| Class 2 | 4 ± 3.1 | **9.5 ± 8** | 4.9 ± 3 | 7.1 ± 4.6 |
| Class 3 | 0.1 ± 0.3 | 0.3 ± 0.6 | 0.2 ± 0.4 | 0.3 ± 0.7 |
| Class 4 | 1 ± 1 | 1.1 ± 1.3 | 1.1 ± 1 | 1.1 ± 0.8 |
| Class 5 | 11.9 ± 4.1 | 12.4 ± 5 | 10.8 ± 4.1 | 13.6 ± 5 |
| *Duration (s)* |  |  |  |  |
| Class 1 | 5.3 ± 1.2 | 3.6 ± 1.4 | 6.2 ± 2.6 | 5.1 ± 2.7 |
| Class 2 | 9.7 ± 4.5 | 7.9 ± 6.4 | 12.5 ± 4 | 8.7 ± 3.9 |
| Class 3 | 1.8 ± 4.6 | 1.6 ± 3.6 | 2.6 ± 5.7 | 2 ± 4.2 |
| Class 4 | 13 ± 12.2 | 9.8 ± 9.8 | **21.2 ± 17.9** | 14.4 ± 10.8 |
| Class 5 | 22.4 ± 10.5 | 16.8 ± 9.1 | **24.9 ± 9** | 17.9 ± 4.7 |
| *Peak frequency (Hz)* |  |  |  |  |
| Class 1 | 1.6 ± 0.5 | 1.9 ± 0.7 | 1.5 ± 0.8 | 1.7 ± 0.8 |
| Class 2 | 0.8 ± 0.6 | 1.3 ± 0.7 | 0.9 ± 0.7 | 1.3 ± 0.6 |
| Class 3 | 0.4 ± 1.4 | 0.3 ± 0.6 | 0.04 ± 0.1 | 0.3 ± 0.7 |
| Class 4 | 0.2 ± 0.2 | 0.3 ± 0.4 | 0.1 ± 0.1 | 0.3 ± 0.2 |
| Class 5 | 0.3 ± 0.1 | 0.4 ± 0.1 | 0.2 ± 0.1 | 0.3 ± 0.1 |
| *Bandwidth* |  |  |  |  |
| Class 1 | 8.6 ± 0.3 | 7.8 ± 2.3 | 8.1 ± 2.2 | 8 ± 2 |
| Class 2 | 8.4 ± 2.7 | 8.4 ± 2.4 | 9.3 ± 0.4 | 9 ± 0.4 |
| Class 3 | 1.4 ± 3.4 | 1.9 ± 3.8 | 1.8 ± 3.9 | 2 ± 4 |
| Class 4 | 5.6 ± 5 | 5.5 ± 4.9 | 6.7 ± 4.7 | 7.5 ± 4.1 |
| Class 5 | 9.6 ± 0.1 | 9.5 ± 0.1 | 9.7 ± 0.1 | 9.6 ± 0.1 |
| *Peak power* |  |  |  |  |
| Class 1 | 6.9 ± 2.4 | 11 ± 6.4 | 9.3 ± 10.7 | 9.4 ± 7.7 |
| Class 2 | 17.4 ± 23.1 | 16 ± 10.8 | 16.1 ± 13.1 | 15 ± 12 |
| Class 3 | 0.9 ± 2.8 | 2.5 ± 5.9 | 1.6 ± 3.5 | 2.2 ± 5.7 |
| Class 4 | 4.4 ± 4.2 | 7.2 ± 7 | 14.4 ± 25.1 | 10.3 ± 9.5 |
| Class 5 | 12.1 ± 3.2 | 17.7 ± 6.7 | 19.7 ± 18.4 | 18.3 ± 13.1 |
| *Max amplitude* |  |  |  |  |
| Class 1 | 207.3 ± 105.2 | 237.9 ± 156 | 150.1 ± 76.8 | 250.5 ± 147.1 |
| Class 2 | 431.4 ± 252.1 | 354.3 ± 223.8 | 290.2 ± 137.9 | 471.5 ± 255.7 |
| Class 3 | 121.9 ± 337.7 | 81.3 ± 171.1 | 70.1 ± 153.2 | 102.8 ± 228.6 |
| Class 4 | 447 ± 415.2 | 296.6 ± 319.7 | 286.5 ± 278.5 | **575.3 ± 451.5** |
| Class 5 | 422.1 ± 221.9 | 426.9 ± 245.9 | 296.5 ± 150 | 466.3 ± 246 |
| *Avg Amplitude* |  |  |  |  |
| Class 1 | 92.7 ± 50.3 | 115.7 ± 73.6 | 68.1 ± 31.2 | 115.9 ± 70.2 |
| Class 2 | 149.6 ± 89.8 | 153.7 ± 102.7 | 106.9 ± 41.2 | 183.4 ± 104.3 |
| Class 3 | 31.8 ± 79.9 | 22 ± 46.3 | 18 ± 38.7 | 30 ± 71 |
| Class 4 | 167.2 ± 171.4 | 129 ± 139.6 | 117.5 ± 114.1 | **227 ± 208.3** |
| Class 5 | 174.4 ± 93.2 | 200 ± 123.9 | 126.4 ± 60.7 | 207.9 ± 122.9 |
| *Max amplitude (%)* |  |  |  |  |
| Class 1 | 30.2 ± 7.8 | 34.6 ± 12.2 | 32.4 ± 9.6 | 33.5 ± 9.8 |
| Class 2 | 58 ± 18.9 | 59.3 ± 19.5 | 63.8 ± 7.6 | 62.5 ± 5.3 |
| Class 3 | 14.9 ± 35.1 | 18.5 ± 37 | 17.6 ± 38 | 19.1 ± 37.3 |
| Class 4 | 51 ± 45.6 | 51.9 ± 47.1 | 65 ± 45.4 | 71 ± 39.8 |
| Class 5 | 60.1 ± 8.4 | 64.4 ± 11.5 | 64.4 ± 8.6 | 61.9 ± 6.7 |
| *Avg Amplitude (%)* |  |  |  |  |
| Class 1 | 13.3 ± 3.4 | 17.3 ± 6.8 | 15.2 ± 4.8 | 15.7 ± 5.2 |
| Class 2 | 19.9 ± 7.2 | 25.8 ± 8.7 | 24.6 ± 5.3 | 24 ± 4 |
| Class 3 | 3.3 ± 8.1 | 5.2 ± 10.7 | 4.6 ± 9.8 | 5.4 ± 11 |
| Class 4 | 18.4 ± 17.4 | 23.7 ± 22.8 | 27 ± 21.3 | 27.1 ± 16.9 |
| Class 5 | 24.1 ± 5.4 | 30.8 ± 7.4 | 27.9 ± 4.2 | 27.3 ± 5.5 |

Data represents mean ± SD.

Additionally, female *Fmr1* KO mice exhibited more small episodes (Class 1) than male FVB-WT mice. Episode duration was altered in *Fmr1* KO mice, particularly in males (Figure 4Bii, Table 3, Supp. Table 1). Multi-burst, rhythmic episodes (Class 4) were shorter in male *Fmr1* KO mice compared to male FVB-WT, and shorter in both female groups (*Fmr1* KO and FVB-WT) compared to male FVB-WT. Similarly, multi-burst, non-rhythmic episodes (Class 5) were shorter in both female and male *Fmr1* KO mice compared to male FVB-WT (Figure 4Bii, Table 3, Supp. Table 1).

Episode amplitude was selectively increased in male *Fmr1* KO mice. Both maximum and average amplitudes of multi-burst, rhythmic episodes (Class 4) were significantly larger in male *Fmr1* KO compared to male FVB-WT and female *Fmr1* KO mice (Figure 4Bvii, Table 3, Supp. Table 1).

Fictive locomotor activity in the *Fmr1* KO mice during early development is similar to that observed in control mice, independently of sex.

We focused on discerning variations in locomotor activity patterns between *Fmr1* KO and FVB-WT mice during early developmental stages (P3-P5). To achieve this, we used an isolated *in vitro* spinal cord preparation (Figure 5A) to record the neuronal activity from the ventral roots at the lumbar L2 and L5 segments in both FVB-WT and *Fmr1* KO mice ^25^. Fictive locomotion was evoked by bath applying N-methyl-D-aspartate (NMDA), serotonin (5-HT), and dopamine (DA). After bath application of the drugs, we observed a robust and coordinated pattern of locomotor-like activity in both groups that stabilized in under 30 minutes (Figure 5B).

**Figure 5:**
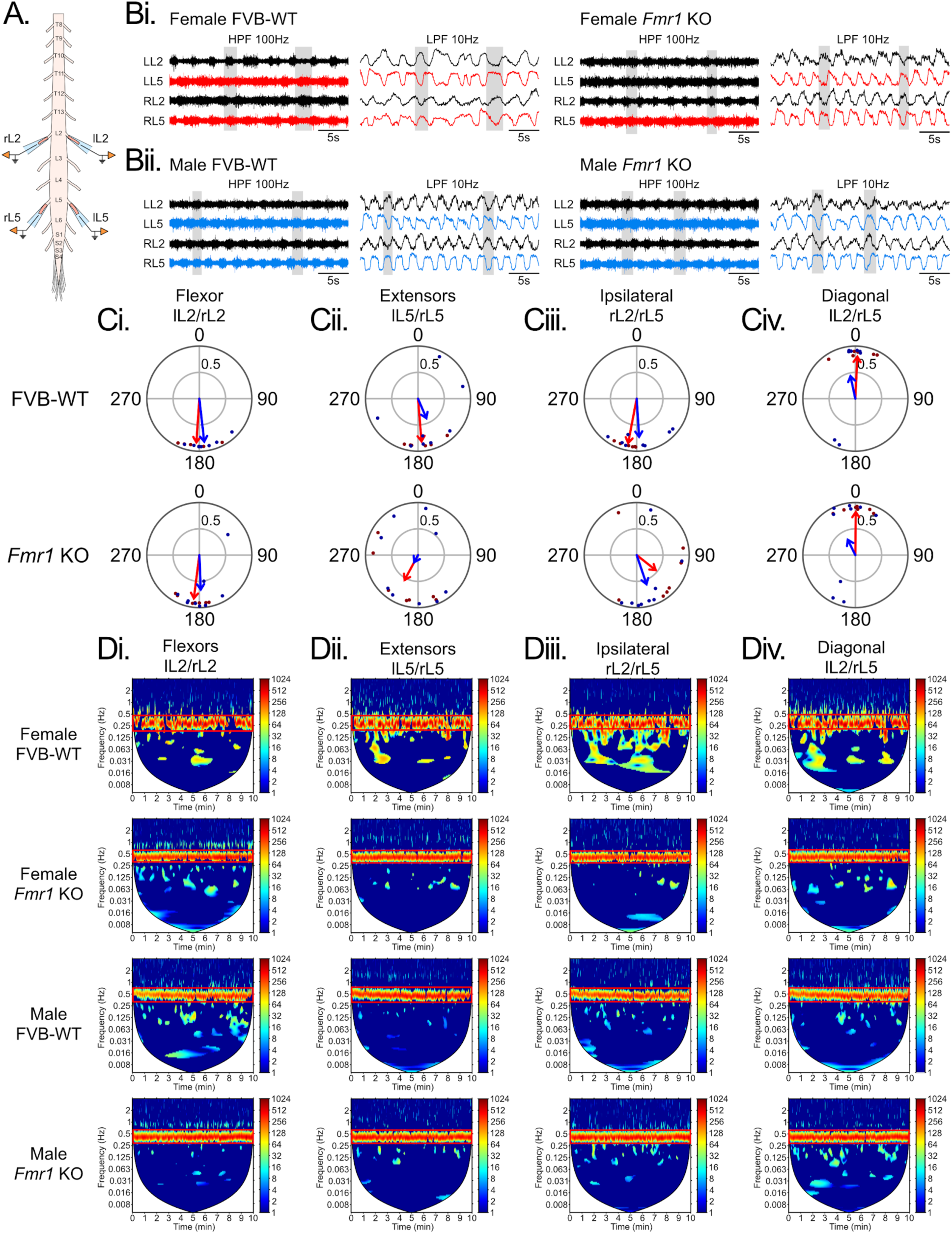
Fictive locomotor activity in the neonate *Fmr1* KO mice is not different from FVB-WT control mice during early development. **A.** Schematic representation of the isolated mouse spinal cord *in vitro* preparation and the ventral root recording electrodes configuration. Suction electrodes were placed in both left (l) and right (r) ventral roots at the lumbar (L2) and (L5) segments. **B.** Raw ventral root neurograms recorded during drug-evoked fictive locomotor activity in female (Bi) and male (Bii) FVB-WT and *Fmr1* KO mice. Rectified and integrated traces are presented in the right panels. Ipsilateral lumbar L2 (black traces) and L5 (blue traces) segments show alternating activity, same for left/right neurograms. **C.** Phase analysis of the interlimb coordination between flexor lL2/rL2 (Ci), extensor lL5/rL5 (Cii), ipsilateral rL2/rL5 (Ciii) and diagonal lL2/rL5 (Civ) limbs for female and male FVB and *Fmr1* KO mice. Phase values of 180° correspond to strict alternation, whereas phase values of 0° correspond to strict synchrony. The mean vector is shown as a red (females) or blue (males) arrow, where arrow direction represents the mean phase relationship and arrow length (mean vector length) reflects how tightly phase values are clustered. Individual phase values are shown as dark red or dark blue dots, respectively. The inner circle indicates a significance level of *p* = 0.05. **D.** Spectral analyses were used to measure the frequency power of the interlimb coordination between flexor lL2/rL2 (Di), extensor lL5/rL5 (Dii), ipsilateral rL2/rL5 (Diii) and contralateral lL2/rL5 (diagonal, Div) limbs in female and male FVB-WT and *Fmr1* KO mice, respectively. Red boxes highlight the analyzed frequency band.

Left-right flexor coordination at L2 was highly consistent across all groups. Both female and male FVB-WT and *Fmr1* KO mice showed strong bilateral alternation of flexor activity with phases clustering tightly around 180° (Figure 5Ci). The high r values (0.8-0.9 - designated by arrows in Figure 5C) indicate consistent phase coupling across trials or cycles, with all groups showing significant phase clustering, demonstrating that basic left-right alternating flexor coordination is preserved in the Fragile X model. Extensor coordination at L5 showed more variability, particularly in males (Figure 5Cii, Table 4). While female mice maintained moderate phase clustering in bilateral extensor alternation (r = 0.8 in FVB-WT, though *Fmr1* KO females show a phase shift to 208°); on the other hand, male mice showed much weaker coupling. The male *Fmr1* KO group showed low phase clustering (r = 0.2) which was not significant (*p* = 0.6310), suggesting variable extensor coordination and a potential sex-specific deficit in lumbar extensor circuit function (Figure 5Ci, Cii, Table 4).

**Table 4.** Interlimb coordination analysis of the neonate fictive locomotion in FVB-WT and *Fmr1* KO mice.

|  | Mean angle | Mean vector length (r) | Z-stat | p |
| --- | --- | --- | --- | --- |
| **Female FVB-WT (n = 6)** |  |  |  |  |
| Flexor IL2/rL2 | 183.9 ± 17.6 | 0.8685 | 5.4584 | 0.0043 |
| Extensor IL5/rL5 | 175.4 ± 18.7 | 0.8496 | 5.392 | 0.0046 |
| Ipsilateral rL2/rL5 | 190.9 ± 9.9 | 0.8961 | 5.8223 | 0.0030 |
| Diagonal IL2/rL5 | 2.8 ± 20.1 | 0.8338 | 5.2978 | 0.0050 |
| **Female *Fmr1* KO (n = 7)** |  |  |  |  |
| Flexor IL2/rL2 | 187.3 ± 15.4 | 0.8651 | 6.5051 | 0.0015 |
| Extensor IL5/rL5 | 208.4 ± 52.1 | 0.5756 | 3.0611 | 0.0468 |
| Ipsilateral rL2/rL5 | 128.4 ± 61.9 | 0.4975 | 2.1807 | 0.1130 |
| Diagonal IL2/rL5 | 0.6 ± 16.4 | 0.8545 | 6.4483 | 0.0016 |
| **Male FVB-WT (n = 8)** |  |  |  |  |
| Flexor IL2/rL2 | 173.5 ± 17.9 | 0.8572 | 7.2565 | 0.0007 |
| Extensor IL5/rL5 | 156.9 ± 65.8 | 0.4411 | 2.131 | 0.1187 |
| Ipsilateral rL2/rL5 | 176.4 ± 29.3 | 0.7624 | 6.1209 | 0.0022 |
| Diagonal IL2/rL5 | 347.9 ± 64.8 | 0.4748 | 2.2285 | 0.1077 |
| **Male *Fmr1* KO (n = 9)** |  |  |  |  |
| Flexor IL2/rL2 | 177.6 ± 17.9 | 0.6918 | 5.9171 | 0.0027 |
| Extensor IL5/rL5 | 208 ± 98.5 | 0.1916 | 0.4604 | 0.6310 |
| Ipsilateral rL2/rL5 | 161.4 ± 48.8 | 0.6208 | 4.3328 | 0.0131 |
| Diagonal IL2/rL5 | 333.8 ± 75.8 | 0.3627 | 1.5419 | 0.2140 |

Data represents mean ± SD.

Flexor-extensor coordination patterns were generally maintained, but with varying strength. Ipsilateral flexor-extensor neurogram pairs showed alternation near 180° in most groups, though with weakened coupling in some cases (particularly female *Fmr1* KO, where coupling became non-significant, *p* = 0.1130, Figure 5Ciii, Table 4). Contralateral flexor-extensor neurogram pairs showed appropriate in-phase bursts for female FVB-WT and *Fmr1* KO mice (near 0°), though male *Fmr1* KO mice showed weak, non-significant phase clustering (Figure 5Civ, Table 4).

There were no significant differences between *Fmr1* KO and FVB-WT mice within each sex (Supplementary Table 1), though the trend toward weaker coordination in male *Fmr1* KO extensors and altered phase relationships in some groups suggests subtle circuit-level changes. Further cross-wavelet analysis showed a strong rhythmicity between segmental, ipsilateral and diagonal ventral root neurograms (Figure 5D, Table 5). Both FVB-WT and *Fmr1* KO male and female mice exhibited this distinct pattern, indicating that the basic spinal circuit for rhythmic locomotor generation is functional in FVB-WT and *Fmr1* KO mice during early development.

**Table 5.** Cross-wavelet analysis of the interlimb coordination in the neonate fictive locomotion.

|  | Mean frequency | Mean coherence | Mean Power |
| --- | --- | --- | --- |
| **Female FVB-WT (n = 6)** |  |  |  |
| Flexor IL2/rL2 | 4.2 ± 0.1 | 1 ± 0.04 | 160.1 ± 41.6 |
| Extensor IL5/rL5 | 0.4 ± 0.1 | 0.9 ± 0.03 | 133.3 ± 21.3 |
| Ipsilateral rL2/rL5 | 0.4 ± 0.1 | 0.9 ± 0.03 | 134.3 ± 31.8 |
| Diagonal IL2/rL5 | 0.4 ± 0.1 | 0.9 ± 0.02 | 139.1 ± 18.4 |
| **Female *Fmr1* KO (n = 7)** |  |  |  |
| Flexor IL2/rL2 | 0.5 ± 0.1 | 0.8 ± 0.1 | 120.8 ± 55.2 |
| Extensor IL5/rL5 | 0.5 ± 0.1 | 0.8 ± 0.04 | 102.4 ± 34.9 |
| Ipsilateral rL2/rL5 | 0.4 ± 0.1 | 0.8 ± 0.04 | 107.6 ± 40.1 |
| Diagonal IL2/rL5 | 0.5 ± 0.1 | 0.9 ± 0.05 | 106 ± 49.7 |
| **Male FVB-WT (n = 8)** |  |  |  |
| Flexor IL2/rL2 | 0.5 ± 0.1 | 0.9 ± 0.04 | 131.2 ± 32.3 |
| Extensor IL5/rL5 | 0.5 ± 0.1 | 0.9 ± 0.04 | 122.1 ± 23.2 |
| Ipsilateral rL2/rL5 | 0.5 ± 0.04 | 0.9 ± 0.05 | 107.9 ± 22.3 |
| Diagonal IL2/rL5 | 0.5 ± 0.1 | 0.9 ± 0.03 | 119 ± 19.2 |
| **Male *Fmr1* KO (n = 9)** |  |  |  |
| Flexor IL2/rL2 | 0.4 ± 0.2 | 0.9 ± 0.1 | 168.8 ± 65.5 |
| Extensor IL5/rL5 | 0.4 ± 0.2 | 0.9 ± 0.1 | 146.2 ± 57.3 |
| Ipsilateral rL2/rL5 | 0.4 ± 0.1 | 0.9 ± 0.1 | 138.4 ± 53.5 |
| Diagonal IL2/rL5 | 0.4 ± 0.2 | 0.9 ± 0.1 | 161.2 ± 49.6 |

Data represents mean ± SD.

### Locomotor activity in the adult *Fmr1* KO mice is comparable to that observed in control mice independently of sex

We next explored the motor deficits in adult *Fmr1* KO and FVB-WT mice. We examined locomotor activity during unmonitored tasks by using the open field test (OFT) and a walkway apparatus for both groups. After 30 minutes in the OFT, the trajectory-tracking of the tested groups (Figure 6A) revealed no preference for activity in the center of the OFT compared to thigmotactic behavior. Further analysis showed an increase in the total distance (Figure 6Bi, Table 6, Supp. Table 1), average speed (Figure 6Bii, Table 6, Supp. Table 1) and max speed (Figure 6Biii, Table 6, Supp. Table 1) for the male *Fmr1* KO mice compared to male FVB-WT mice, and only an increase in the max speed for the female *Fmr1* KO mice compared to female FVB-WT mice (Figure 6Biii, Supp. Table 1). These results replicated previous work showing hyperactivity patterns in male *Fmr1* KO mice ^16,17^.

**Figure 6:**
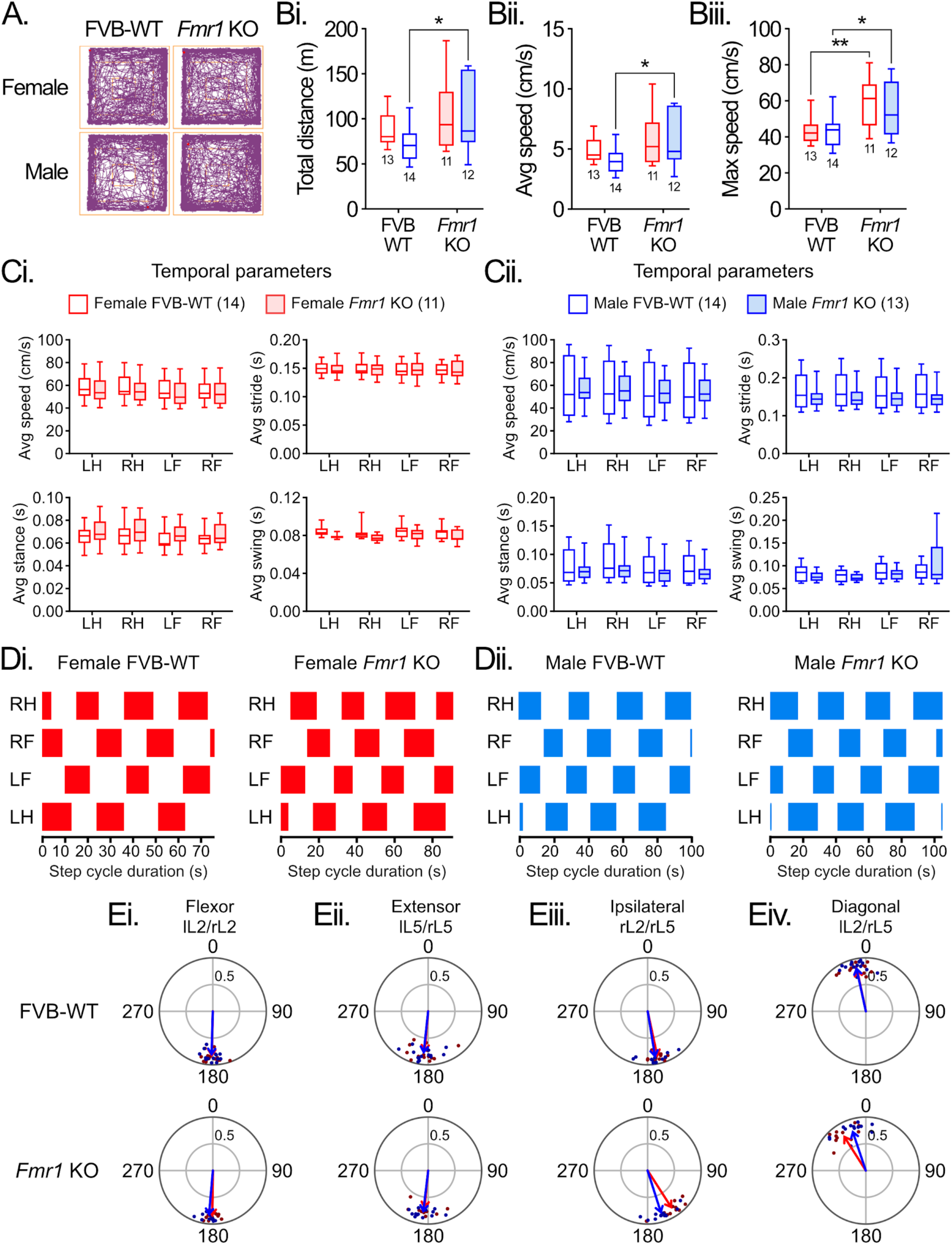
Adult male *Fmr1* KO mice are hyperactive compared to control FVB-WT mice, with no differences in their locomotor pattern. **A.** Representative trace of the locomotor trajectory of the mice during OFT in both male and female FVB-WT and *Fmr1* KO mice. **B.** Box and whisker plots display interquartile range (boxes), median (horizontal lines), maximum, and minimum values in data range (whiskers) for the analysis of the OFT. Total distance (Bi), average (Bii) and max speeds (Biii) were significantly increased in the male *Fmr1* KO compared to the control FVB-WT mice. The female group showed a higher maximum speed in the *Fmr1* KO than in the FVB-WT mice. **C.** Similar to B, for the analysis of the temporal parameters of gait during the walkway test. No differences were observed between the *Fmr1* KO and FVB-WT mice in the female (Ci) or male (Cii) groups. **D.** Representative gait cycle diagrams for the *Fmr1* KO and FVB-WT, female (Di) and male (Dii) mice. **E.** Phase analysis of the interlimb coordination between forelimbs (Ei), hindlimbs (Eii), ipsilateral (Eiii) and contralateral (diagonal, Eiv) limbs for the *Fmr1* KO and FVB-WT, female and male mice. Phase values of 180° correspond to strict alternation, whereas phase values of 0° correspond to strict synchrony. The mean vector is represented as a red (females) or blue (males) arrow, single values are shown as dark red or dark blue dots, respectively. The inner circle indicates a significance level of *p* = 0.05. Two-way ANOVA followed by Sidák multiple comparison test (*: *p* < 0.05, **: *p* < 0.01, ***: *p* < 0.001). LH, left hindlimb; LF, left forelimb; RH, right hindlimb; RF, right forelimb.

**Table 6.** Open field test analysis of adult FVB-WT and *Fmr1* KO mice.

|  | ♀ FVB (n = 13) | ♀ Fmr1 KO (n = 11) | ♂ FVB (n = 14) | ♂ Fmr1 KO (n = 12) |
| --- | --- | --- | --- | --- |
| Total distance (m) | 88.5 ± 18.1 | 106.9 ± 41.7 | 71.7 ± 18.7 | 102.6 ± 41.5 |
| Average speed (cm/s) | 4.9 ± 1 | 6 ± 2.3 | 4 ± 1 | 5.7 ± 2.3 |
| Max speed (cm/s) | 42.8 ± 6.7 | 58.8 ± 12.9 | 43.9 ± 9.4 | 55 ± 15.6 |

Data represents mean ± SD.

Next, we recorded both groups of mice during freely moving activity using our walkway apparatus ^35^, and semi-automatic analysis was performed using our Visual Gait Lab suite ^36^. Basic temporal gait parameters were normal in *Fmr1* KO mice of both sexes. Neither female nor male *Fmr1* KO mice showed significant differences from wild-type controls in walking speed, stride duration, stance duration, or swing duration across all limbs (Figure 6Ci, Table 7, Supp. Table 1). These findings indicate that the fundamental timing of individual limb movements during overground locomotion is preserved in the Fragile X model. Figure 6D shows the locomotor pattern of activity in the female (red) and male (blue) *Fmr1* KO and FVB-WT mice.

**Table 7.** Temporal parameters quantified during the Walkway test in adult FVB-WT and *Fmr1* KO mice.

|  | ♀ FVB (n = 14) | ♀ Fmr1 KO (n = 11) | ♂ FVB (n = 14) | ♂ Fmr1 KO (n = 13) |
| --- | --- | --- | --- | --- |
| *Average speed (cm/s)* |  |  |  |  |
| LH | 57.9 ± 10.7 | 55.9 ± 12.7 | 56.6 ± 25.5 | 56.8 ± 14.4 |
| RH | 58.3 ± 10.9 | 55.7 ± 12.4 | 55.7 ± 23.8 | 57.2 ± 15.2 |
| LF | 55.2 ± 10.5 | 53.4 ± 11.8 | 54 ± 23.6 | 54.1 ± 14.4 |
| RF | 55.2 ± 10.3 | 53.5 ± 11.7 | 53.9 ± 23.7 | 54.6 ± 13.6 |
| *Average stride (s)* |  |  |  |  |
| LH | 0.150 ± 0.011 | 0.148 ± 0.014 | 0.165 ± 0.048 | 0.149 ± 0.027 |
| RH | 0.149 ± 0.014 | 0.148 ± 0.014 | 0.167 ± 0.046 | 0.148 ± 0.027 |
| LF | 0.147 ± 0.014 | 0.147 ± 0.016 | 0.164 ± 0.047 | 0.149 ± 0.030 |
| RF | 0.146 ± 0.013 | 0.147 ± 0.015 | 0.165 ± 0.046 | 0.147 ± 0.027 |
| *Average stance (s)* |  |  |  |  |
| LH | 0.065 ± 0.009 | 0.069 ± 0.013 | 0.081 ± 0.031 | 0.071 ± 0.018 |
| RH | 0.067 ± 0.011 | 0.07 ± 0.012 | 0.088 ± 0.034 | 0.074 ± 0.020 |
| LF | 0.063 ± 0.009 | 0.066 ± 0.010 | 0.077 ± 0.028 | 0.067 ± 0.018 |
| RF | 0.063 ± 0.008 | 0.068 ± 0.011 | 0.077 ± 0.028 | 0.068 ± 0.016 |
| *Average swing (s)* |  |  |  |  |
| LH | 0.084 ± 0.005 | 0.079 ± 0.002 | 0.084 ± 0.018 | 0.077 ± 0.011 |
| RH | 0.082 ± 0.007 | 0.077 ± 0.004 | 0.079 ± 0.014 | 0.074 ± 0.008 |
| LF | 0.084 ± 0.007 | 0.081 ± 0.006 | 0.088 ± 0.020 | 0.082 ± 0.013 |
| RF | 0.083 ± 0.007 | 0.079 ± 0.007 | 0.088 ± 0.020 | 0.104 ± 0.049 |

Data represents mean ± SD.

Inter-limb coordination patterns were intact in *Fmr1* KO mice. Phase analysis revealed the expected quadrupedal gait pattern in all groups: homologous limbs (left-right forelimbs and hindlimbs) alternated appropriately, ipsilateral limbs (forelimb-hindlimb on the same side) showed proper alternation, and diagonal limbs moved in-phase as expected during normal walking (Figure 6Ei-ii, Table 8, Supp. Table 1). No significant differences between *Fmr1* KO and wild-type mice in any of these coordination measures for either females or males were found (Supp. Table 1), demonstrating that the basic quadrupedal locomotor coordination network functions normally in the Fragile X model.

**Table 8.**
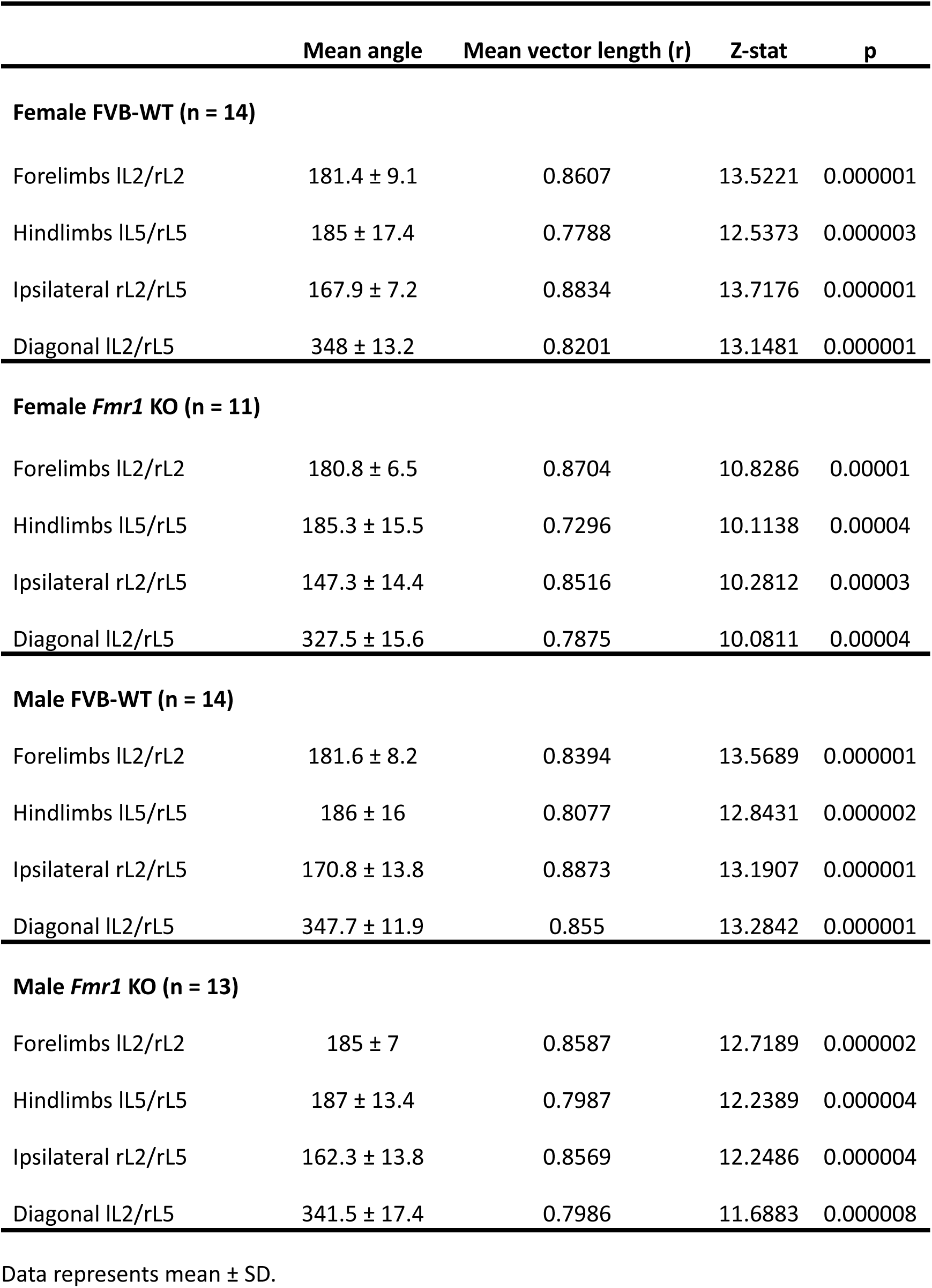
Interlimb coordination analysis of the adult locomotor activity in FVB-WT and *Fmr1* KO mice.

### *Fmr1* KO mice have a low performance during skill-demanding tasks compared to FVB-WT in both female and male mice

To analyze the ability of the *Fmr1* KO mice in a more skilled-demanding task, we performed a ladder rung test (LRT) using two different patterns for the rung placement, a regularly spaced and an irregular pattern (patterns A and B, respectively, Figure 7A) ^37,38^ to prevent the mice from memorizing it.

**Figure 7:**
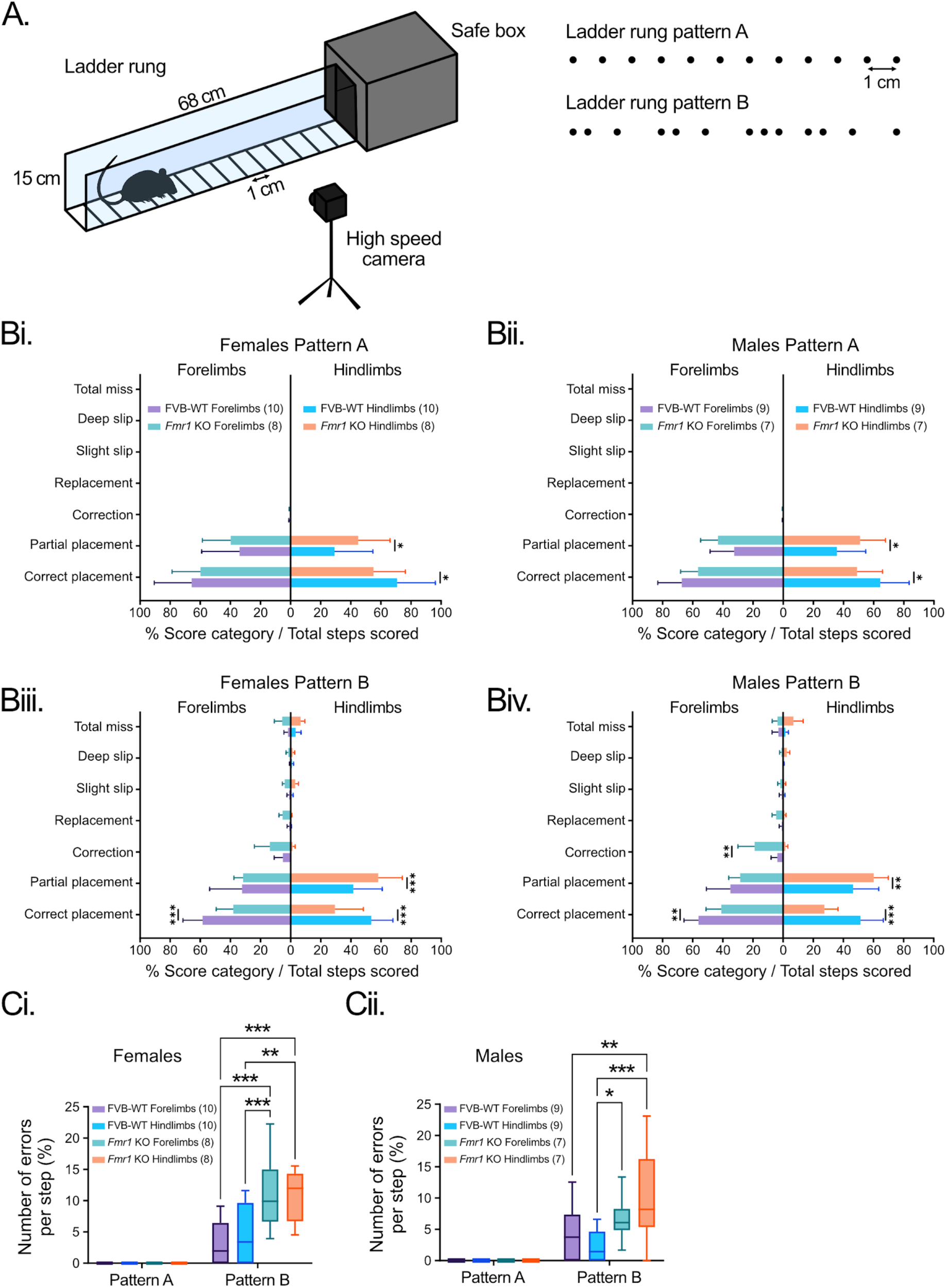
*Fmr1* KO mice showed difficulties when performing a skill-demanding walking task compared to FVB-WT mice. **A.** Schematic of the ladder rung apparatus used. The rung separation was 1 cm for pattern A, with a regularly spaced pattern; pattern B used an irregularly spaced distribution of rungs. Mice were tracked using a fixed high-speed camera. **B.** Box and whisker plots display interquartile range (boxes), median (horizontal lines), maximum, and minimum values in data range (whiskers) for the analysis of the 7 categories scored from recorded videos during tasks using both pattern A (Bi, Bii) and B (Biii, Biv). Forelimbs and hindlimbs from the same mice were analyzed during patterns A and B. **C.** Analysis of the number of errors per step in female (Ci) and male (Cii) FVB-WT and *Fmr1* KO mice. Two-way ANOVA followed by Sidák multiple comparison test (*: *p* < 0.05, **: *p* < 0.01, ***: *p* < 0.001).

**Figure 8:**
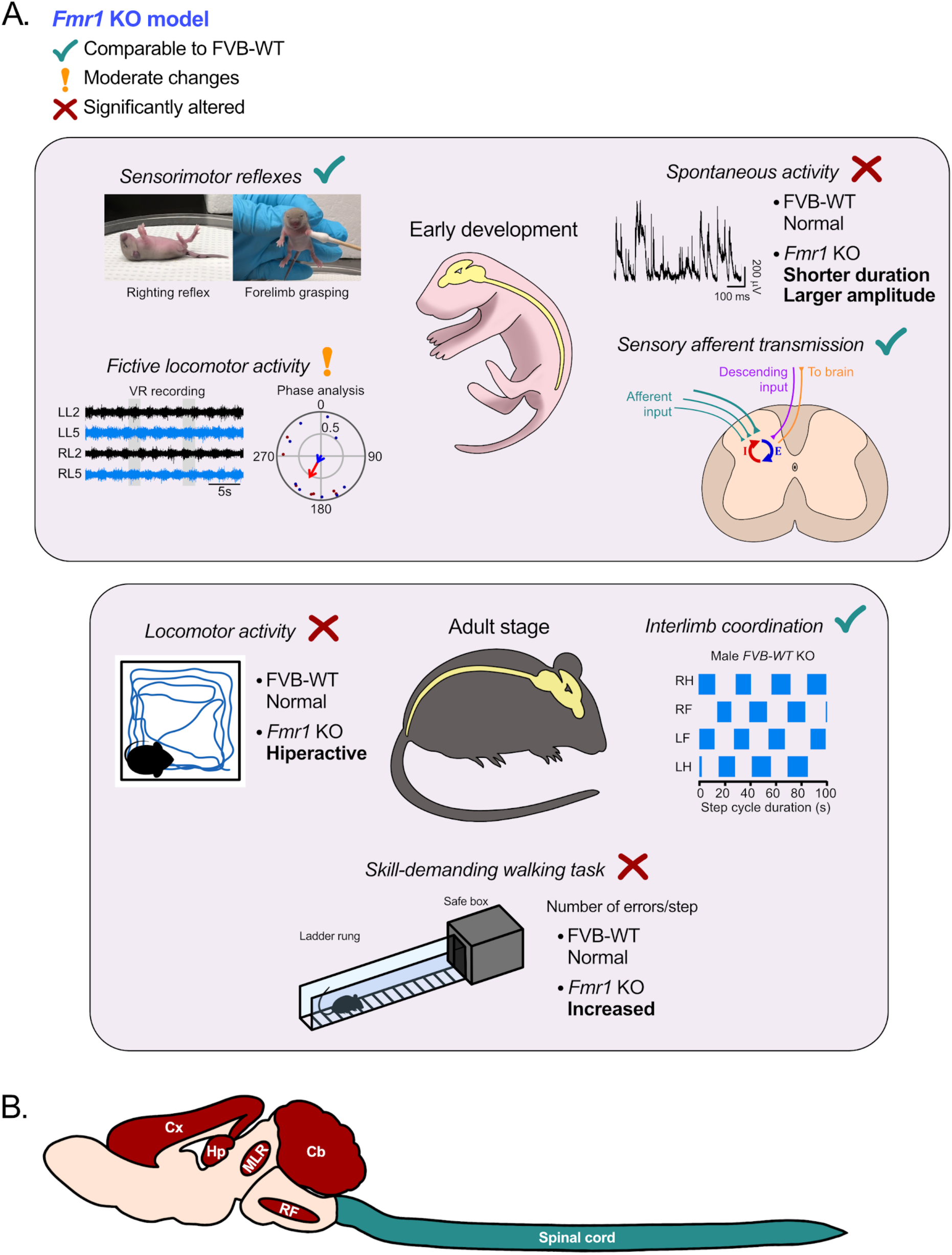
Summary of the sensorimotor development differences in the *Fmr1* KO compared to FVB-WT during early development and adult stage. **A.** Schematic depicting the sensorimotor development differences between *Fmr1* KO and FVB-WT mice during early development (top panel) and in the adult stage (bottom panel). Teal color checkmarks denote comparable results for the *Fmr1* KO mice compared to FVB-WT. Orange exclamation marks highlight modest changes in the *Fmr1* KO, whereas red crosses indicate significant differences compared to the FVB-WT controls. **B.** Depicts the proposed supraspinal areas implicated in the *Fmr1*-mediated dysfunction. Since major dysfunction is observed during skill-demanding walking tasks in the adult *Fmr1* KO mice, the spinal cord is suggested not to be involved. Cx, Cortex; Hp, Hippocampus; MLR, Mesencephalic locomotor region; RF, Reticular formation; Cb, Cerebellum.

Using the regularly spaced rungs from pattern A in the LRT, we observed a reduction in the number of correct placements and an increase in the partial placements of the hindlimbs for the *Fmr1* KO compared to FVB-WT mice, in both female and male groups (Figure 7Bi-ii, Table 9, Supp. Table 1). Analysis of the performance during the irregularly spaced pattern B showed a reduction in the number of correct placements in both forelimbs and hindlimbs for female and male groups (Figure 7Biii-iv, Table 9, Supp. Table 1). The male group also showed an increased number of partial placements for the *Fmr1* KO compared to FVB-WT (Figure 7Biv, Table 9, Supp. Table 1). We observed a reduction in the number of correct placements and an increase in partial hindlimb placements in *Fmr1* KO compared to FVB-WT mice, in both female and male groups (Figure 7Biii-iv, Table 9, Supp. Table 1).

**Table 9.**
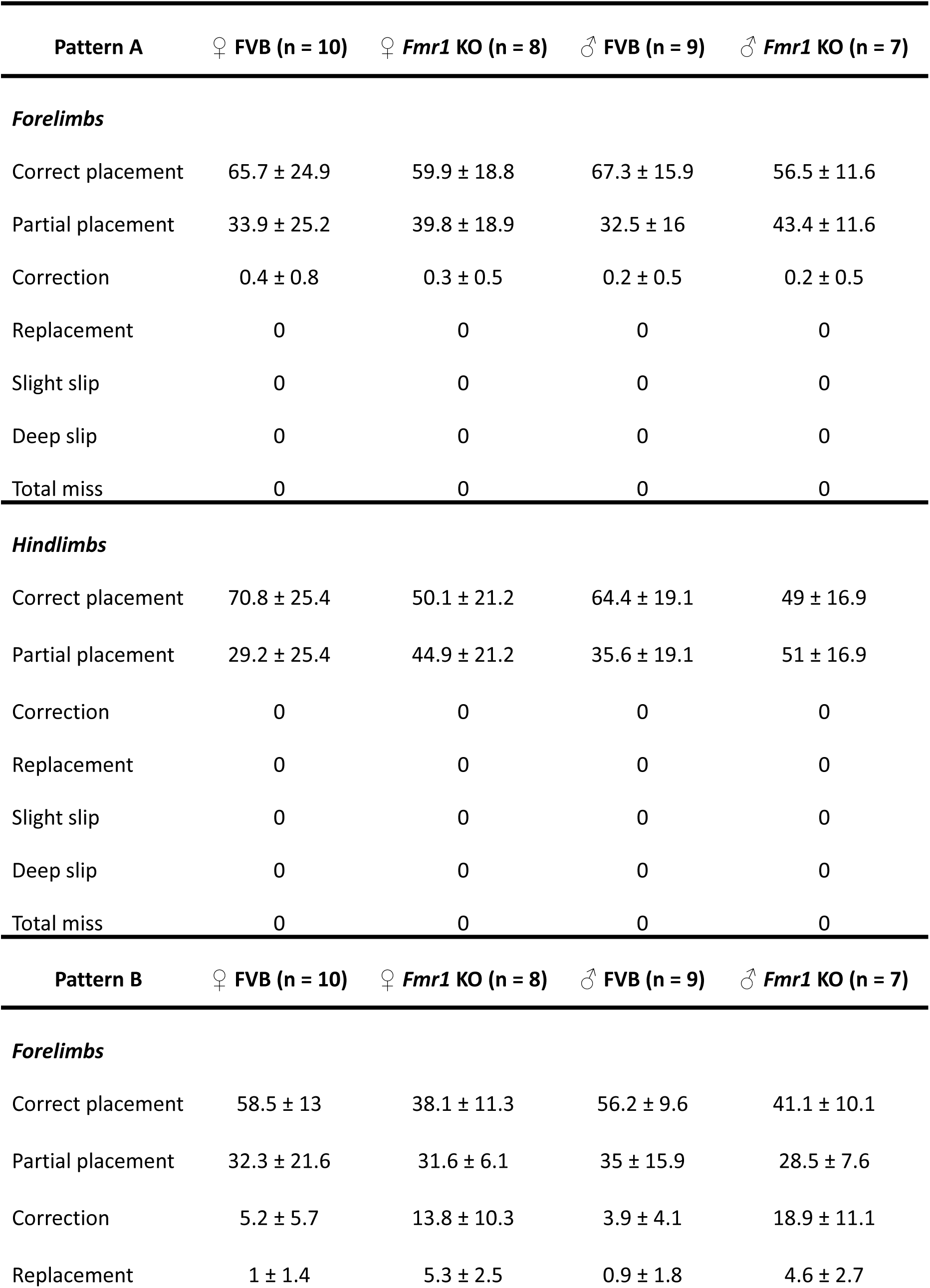
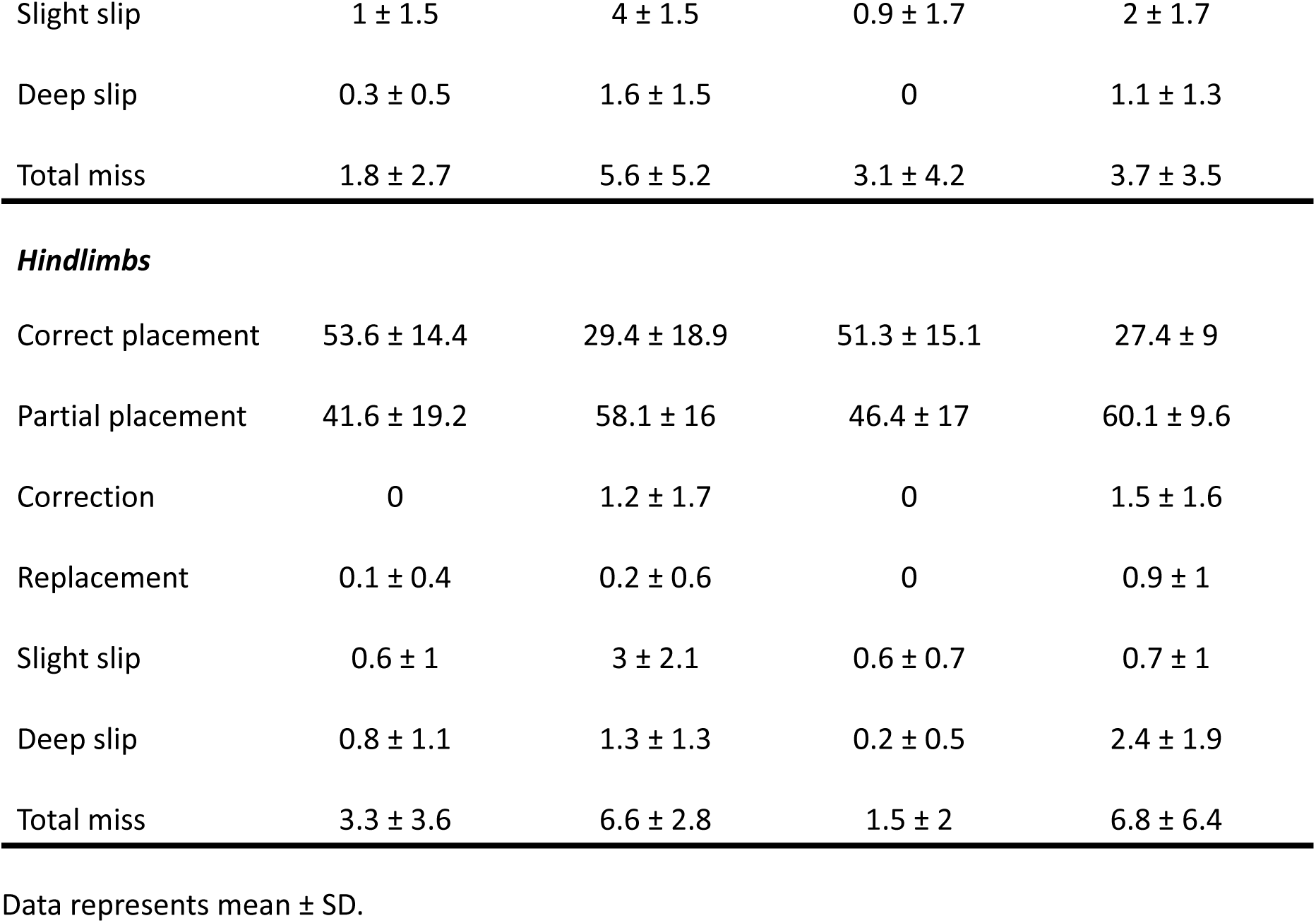
Ladder rung scoring during patterns A and B, in the FVB-WT and *Fmr1* KO mice.

| Pattern A | ♀ FVB (n = 10) | ♀ *Fmr1* KO (n = 8) | ♂ FVB (n = 9) | ♂ *Fmr1* KO (n = 7) |
| --- | --- | --- | --- | --- |
| ***Forelimbs*** |  |  |  |  |
| Correct placement | 65.7 ± 24.9 | 59.9 ± 18.8 | 67.3 ± 15.9 | 56.5 ± 11.6 |
| Partial placement | 33.9 ± 25.2 | 39.8 ± 18.9 | 32.5 ± 16 | 43.4 ± 11.6 |
| Correction | 0.4 ± 0.8 | 0.3 ± 0.5 | 0.2 ± 0.5 | 0.2 ± 0.5 |
| Replacement | 0 | 0 | 0 | 0 |
| Slight slip | 0 | 0 | 0 | 0 |
| Deep slip | 0 | 0 | 0 | 0 |
| Total miss | 0 | 0 | 0 | 0 |
| ***Hindlimbs*** |  |  |  |  |
| Correct placement | 70.8 ± 25.4 | 50.1 ± 21.2 | 64.4 ± 19.1 | 49 ± 16.9 |
| Partial placement | 29.2 ± 25.4 | 44.9 ± 21.2 | 35.6 ± 19.1 | 51 ± 16.9 |
| Correction | 0 | 0 | 0 | 0 |
| Replacement | 0 | 0 | 0 | 0 |
| Slight slip | 0 | 0 | 0 | 0 |
| Deep slip | 0 | 0 | 0 | 0 |
| Total miss | 0 | 0 | 0 | 0 |
| Pattern B | ♀ FVB (n = 10) | ♀ *Fmr1* KO (n = 8) | ♂ FVB (n = 9) | ♂ *Fmr1* KO (n = 7) |
| ***Forelimbs*** |  |  |  |  |
| Correct placement | 58.5 ± 13 | 38.1 ± 11.3 | 56.2 ± 9.6 | 41.1 ± 10.1 |
| Partial placement | 32.3 ± 21.6 | 31.6 ± 6.1 | 35 ± 15.9 | 28.5 ± 7.6 |
| Correction | 5.2 ± 5.7 | 13.8 ± 10.3 | 3.9 ± 4.1 | 18.9 ± 11.1 |
| Replacement | 1 ± 1.4 | 5.3 ± 2.5 | 0.9 ± 1.8 | 4.6 ± 2.7 |
| Slight slip | 1 ± 1.5 | 4 ± 1.5 | 0.9 ± 1.7 | 2 ± 1.7 |
| Deep slip | 0.3 ± 0.5 | 1.6 ± 1.5 | 0 | 1.1 ± 1.3 |
| Total miss | 1.8 ± 2.7 | 5.6 ± 5.2 | 3.1 ± 4.2 | 3.7 ± 3.5 |
| ***Hindlimbs*** |  |  |  |  |
| Correct placement | 53.6 ± 14.4 | 29.4 ± 18.9 | 51.3 ± 15.1 | 27.4 ± 9 |
| Partial placement | 41.6 ± 19.2 | 58.1 ± 16 | 46.4 ± 17 | 60.1 ± 9.6 |
| Correction | 0 | 1.2 ± 1.7 | 0 | 1.5 ± 1.6 |
| Replacement | 0.1 ± 0.4 | 0.2 ± 0.6 | 0 | 0.9 ± 1 |
| Slight slip | 0.6 ± 1 | 3 ± 2.1 | 0.6 ± 0.7 | 0.7 ± 1 |
| Deep slip | 0.8 ± 1.1 | 1.3 ± 1.3 | 0.2 ± 0.5 | 2.4 ± 1.9 |
| Total miss | 3.3 ± 3.6 | 6.6 ± 2.8 | 1.5 ± 2 | 6.8 ± 6.4 |

Data represents mean ± SD.

Next we compared the number of errors per step during both tested patterns and found an increase in the number of errors per step for the *Fmr1* KO compared to FVB-WT mice in both female (Figure 7Ci, Table 10, Supp. Table 1) and male groups (Figure 7Cii, Table 10, Supp. Table 1). These results suggest that supraspinal or even descending neural pathways involved in fine adjustments of motor control, balance, limb coordination and muscle control could be affected in the *Fmr1* KO, as they showed an increase in the number of error steps during the most demanding task.

**Table 10.** Quantification of the number of errors per step for the FVB-WT and *Fmr1* KO mice, during the ladder rung test.

| Pattern A | ♀ FVB (n = 10) | ♀ *Fmr1* KO (n = 8) | ♂ FVB (n = 9) | ♂ *Fmr1* KO (n = 7) |
| --- | --- | --- | --- | --- |
| ***Errors/step*** |  |  |  |  |
| Forelimbs | 0 | 0 | 0 | 0 |
| Hindlimbs | 0 | 0 | 0 | 0 |
| Pattern B | ♀ FVB (n = 10) | ♀ *Fmr1* KO (n = 8) | ♂ FVB (n = 9) | ♂ *Fmr1* KO (n = 7) |
| ***Errors/step*** |  |  |  |  |
| Forelimbs | 3.1 ± 3.5 | 11.2 ± 5.8 | 4 ± 4.5 | 6.8 ± 3.6 |
| Hindlimbs | 4.7 ± 5.1 | 11 ± 4.1 | 2.2 ± 2.6 | 10 ± 7.6 |

Data represents mean ± SD.

## Discussion

Our work comprehensively analyzes the role of spinal cord circuits in Fmr1 KO mice. We examine the functional output of the CPG, including spontaneous network activity and fictive locomotion. We analyzed developmental behavioral milestones in terms of motor development, and replicated work in the adult to show hyperactivity and increased errors in paw placement - commonly associated with abnormal descending control of gait.

### Preservation of Functional Spinal CPG Output in Fmr1 KO Mice

Gait patterns were conserved during *unskilled* locomotion (unsupervised walkway data) in adult wildtype and *Fmr1* KO mice, consistent with previous runway findings ^20^. This is important since it is recognized that the basic rhythm can be generated by spinal circuits in adult mice ^39,40^. Our work demonstrates that CPG circuits from *isolated spinal cord preparations* were capable of generating robust, coordinated bouts of locomotor-like activity that were similar to wild-type controls, independent of sex. This was in spite of the demonstrated absence of FMRP within the brain and spinal cord of our *Fmr1* KO mice. The CPG can produce coordinated flexor-flexor and extensor-flexor bursts, both ipsilaterally and contralaterally, with periods that are similar to each other.

While the output may be similar, this does not imply that the circuitry and synaptic strengths are conserved ^41^. In fact, the functional connectome underlying locomotion most likely reflects differences in kinematics and muscle fibre composition across individual mice. Individual neuronal elements have been shown to exhibit substantial individual variability in intrinsic properties and synaptic input - and interestingly, connectivity between elements can act to compensate for this variability ^42^. This concept of network degeneracy, in which different combinations of underlying parameters can produce similar functional outputs, is well recognized ^41,42^ and should be considered an underlying mechanism ^43^. Indeed, multiple classes of interneurons within the spinal cord can be genetically dissected with network output being altered but not eliminated ^43–48^. Whether this extends to the forelimb spinal CPG is unknown; however, the forelimb CPG, in isolation from the lumbar segments, is more challenging to resolve ^49^.

Moreover, this preservation of the CPG output in *Fmr1* KO mice was supported by our gait analysis of adult mice. However, skilled gait is crucial for walking adaptability ^38^, requiring highly precise motor coordination between forelimbs and hindlimbs. Indeed, when we increased the task’s complexity using a ladder rung task, *Fmr1* KO mice showed deficits, with increased footfalls observed in both sexes. The ladder rung is sensitive to age-related motor deficits and to different motor dysfunction models ^50–53^. These data are similar to work showing motor dysfunction observed on rotarods and balance beams in *Fmr1* KO mice ^17,20,54,55^. Therefore, *Fmr1* KO mice exhibit deficits in fine motor control, balance, and limb coordination required for skilled walking.

### Altered Properties of Spontaneous Spinal Network Activity

Maintaining CPG output in both the neonatal and adult states naturally raises questions about the integrity of the underlying network. An interesting hypothesis is that spinal networks show inherent compensatory mechanisms that may maintain CPG function for unskilled locomotor tasks in *Fmr1* KO mice. Therefore, this could represent a compensatory mechanism that allows spinal cord functions to continue in the absence of FMRP.

To address this, we examined the function of spontaneous activity patterns from isolated spinal cord preparations ^25,34,56,57^. These patterns are associated with recurrently connected sets of interneurons that generate periodic activity early in development ^58^, leading to stochastic patterns later, once GABAergic neurons become inhibitory ^59^.

We did observe differences in locomotor network function - notably, the female *Fmr1* KO mice showed a greater number of large non-rhythmic episodes and small episodes compared to male wildtype mice. Moreover, male *Fmr1* KO mice showed shorter multi-burst, rhythmic and non-rhythmic episodes compared to male FVB-WT, and an increased maximum and average amplitude for multi-burst, rhythmic episodes. Collectively, this suggests minor changes in output patterns, with larger-amplitude bursts in *Fmr1* KO mice compared to FVB-WT mice, independent of sex.

Work in other brain areas shows similarities, with increases in input resistance observed in the barrel cortex at P10-12 in *Fmr1* KO mice ^60,61^. This is associated with increased spike output. Modelling supported the idea that compensatory effects mitigate rather than exacerbate network dysfunction in the barrel cortex of *Fmr1* KO mice ^61^. In the visual cortex, there is evidence of unbalanced centrally driven patterns preceding eye opening ^62^. Our work showing higher episode amplitudes agrees with findings in the somatosensory cortex, where 3-fold higher neuronal firing rates were observed in *Fmr1* KO mice ^10^. As noted, GABAergic drive starts as excitatory and becomes inhibitory over time. In *Fmr1* KO mice, work shows a delayed switch from depolarizing to hyperpolarizing GABA transmission in the cortex. If this were the case in the spinal cord, it could lead to the increased amplitudes of spontaneous episodes we observed, particularly in male mice.

### Early Sensory and Developmental Reflexes Are Not Affected by Fmr1 Deletion

We selected P10 for *in vitro* experiments based on key developmental milestones. At this age, eyes remain closed (opening occurs between P11 and P14) ^27,63,64^, and the ear twitch reflex is emerging (appears at P7-9) ^26^. In contrast, the rooting reflex is present at P2 and abates during the early neonatal period (Sarkar et al. 2020). While the ear twitch was absent, P10 was within the developmental stage when it could have been missed. Eyes remained closed, and the rooting reflex was absent. Despite this timing, sensorimotor and vestibular systems, assessed through righting, negative geotaxis, forelimb grasping, and cliff aversion tests, showed no differences between *Fmr1* and FVB-WT mice.

Given the spontaneous network activity changes we observed in spinal circuits, we investigated whether sensory afferent transmission and its presynaptic regulation were similarly affected in early development. Using the isolated spinal cord preparation, we examined dorsal root potentials (DRPs), which reflect presynaptic inhibition ^65^. Testing both low and high stimulation intensities to recruit large- and small-diameter afferents (including C fibers) revealed no amplitude differences across conditions, indicating normal perinatal sensory gating at the spinal level. These findings are noteworthy, as alterations in pain processing are observed in adult *Fmr1* KO mice ^22^, suggesting that presynaptic inhibition deficits in mature animals may reflect altered supraspinal descending modulation of dorsal horn circuits rather than intrinsic spinal abnormalities.

### Motor Deficits in Skilled Locomotion Suggest a Supraspinal Locus of Pathology

The primary motor cortex is essential for accurate paw placement during skilled walking ^66,67^, and corticospinal lesions produce footfault patterns similar to those we observed ^53^. In *Fmr1* KO mice, the motor cortex shows circuit-level pathology similar to that reported in sensory cortices, including altered dendritic spine morphology (Selby et al., 2007), enhanced mGluR-LTD (Hays et al., 2011), and an excitatory-inhibitory imbalance ^60^. These cellular abnormalities manifest behaviorally as motor learning deficits (Padmashri et al., 2013), which would impair the experience-dependent refinement needed for skilled tasks.

Beyond direct corticospinal control, the reticular formation integrates cortical commands with spinal CPG output and mediates postural corrections during locomotion ^68–70^. GABA dysfunction in brainstem nuclei, including the reticular formation ^71^, and altered brainstem-cortical connectivity ^72^ in *Fmr1* KO mice suggest that reticulospinal pathways may contribute to impaired balance and coordination during ladder walking that we observed.

The balance demands of irregular ladder patterns implicate vestibulospinal pathways ^73^, though vestibular function in FXS remains largely unexplored. Additionally, although our focus is descending pathways, cerebellar dysfunction could also contribute indirectly ^74^. Notably, cerebellar lesions produce foot-placement errors on irregular surfaces ^75^, findings similar to ours.

Collectively, these data suggest skilled locomotion deficits in *Fmr1* KO mice could arise from dysfunction across multiple descending systems that integrate cortical planning, cerebellar prediction, and brainstem corrections. The spinal CPG, while showing subtle alterations in spontaneous activity patterns, retains its fundamental capacity for rhythmogenesis - perhaps through the network degeneracy and compensatory mechanisms observed in other neural systems affected by FMRP loss ^61^.

## Conclusion

We found that *Fmr1* KO mice retain the capacity of spinal CPGs to produce basic locomotor behavior, independent of sex, which mirrors our *in vivo* work showing similar gait patterns for unskilled locomotion. Interestingly, we were able to distinguish differences in spontaneous network activity, which raises the question of whether spinal cord circuitry may possess an inherent capacity to compensate for the loss of Fmr1 in *Fmr1* KO. Despite normal unskilled locomotion, skilled locomotion was impaired in *Fmr1* KO mice, suggesting a supraspinal locus for *Fmr1*-mediated dysfunction. Our work identifies the descending supraspinal pathway as a potential target.

## Disclosures

## Acknowledgements

We would like to acknowledge support from the CSMOpto core facility, Spinal Core software was a kind gift from Prof A. Lev-Tov (Hebrew University).

## Funding

We acknowledge studentships from Hotchkiss Brain Institute, NSERC Brain Create (JMC), and the Faculty of Veterinary Medicine. This research is supported by grants provided by the Canadian Institute of Health Research (PJW) and an NSERC Discovery grant (PJW).

## Author contributions

Designed research (JMC, PW), performed research (JMC, AM, LM, MAT), analyzed data (JMC), writing manuscript (JMC, PW), edited manuscript (JMC, PW, NC, MAT, AM, LM).

## Conflict of interests

The authors declare no competing financial interests.

## Methods

### Ethical approval & animals

Experiments were performed on wild-type (Jackson Lab stock #004828, FVB.129P2-*Pde6b^+^ Tyr^c-ch^*/AntJ) and *Fmr1* KO (Jackson Lab stock #004624, FVB.129P2-*Pde6b^+^ Tyr^c-ch^ Fmr1^tm1Cgr^*/J) mice, both sexes. A total of 228 mice were used. Mice were genotyped using PCR according to protocols from Jackson Laboratory. Postnatal 3-10 (P3-P10) and adult (P50-P100) mice were used. Mice were housed on a 12:12 light: dark cycle (lights on 07:00 - off 19:00) at a room temperature of 20°C and relative humidity of 34% with food and water access *ad libitum*. All procedures were approved by the University of Calgary Health Sciences Animal Care Committee (AC24-0140).

### Developmental Milestone assessment in neonate mice

Neonatal mice were assessed for the acquisition of early sensorimotor and reflexive behaviors during postnatal development following the battery of behavioral tests by ^28^ as a guideline. Pups were weighed and examined on postnatal day 10 (P10). Behavioral experiments were performed from 09:00-14:00. Pups were individually identified and tail marked, sex was determined by dissecting them at the end of the session. All assessments were conducted in a quiet room maintained at 24–26 °C. Between tests, pups were returned to their home cage and allowed to recover on nesting material warmed with a heat pad. Each task was scored according to predefined criteria (see below).

### Rooting reflex

The rooting reflex was assessed by gently stroking the perioral region (whisker pad and corner of the mouth) using a fine cotton swab. A positive response was defined as a consistent head turn toward the stimulus accompanied by mouth opening or probing movements within three seconds of stimulation. Each side of the face was tested three times.

### Righting reflex

To evaluate postural control, pups were placed gently on their backs on a flat surface. The latency to return to a prone position with all four paws contacting the surface was recorded, with a maximum cutoff time of 30 seconds. Four different trials per animal were averaged. Failure to right themselves within the cutoff time was scored as unsuccessful.

### Forelimb grasping

Forelimb grasping ability was assessed by lightly touching the palmar surface of each forepaw with a cotton swab. A positive response was defined as flexion of the digits around the swab or a sustained grasp lasting at least one second. Each forelimb was tested three times independently.

### Ear twitch reflex

Ear tactile responsiveness was evaluated by lightly touching the external ear (pinna) with a cotton swab. A positive response was defined as a rapid twitch or retraction of the ear or an associated head movement occurring immediately after stimulation. Each ear was examined three times independently.

### Negative geotaxis

Negative geotaxis was assessed by placing the pup head-downward on a ramp inclined at 40°. The latency to rotate the body 180° to face upward was recorded, with a maximum cutoff of 60 seconds. Three different trials per animal were averaged. Failure to complete reorientation within the cutoff time was scored as unsuccessful.

### Cliff aversion reflex

To assess sensorimotor coordination and vestibular function, pups were placed at the edge of a flat surface with the forepaws and snout extending over the edge, a piece of cotton was placed at the bottom as a safety cushion. The latency to completely withdraw from the edge by backward movements or lateral turnings was measured, with a maximum cutoff of 30 seconds. Three different trials per animal were averaged. Failure to completely withdraw within the cutoff time was scored as unsuccessful.

### Air righting reflex

The air righting reflex was evaluated by holding the pup in a supine position approximately 20 cm above a soft padded surface and releasing it gently. A successful response was defined as rotation of the body in midair to land on all four paws. This test was performed once at the end to minimize stress.

### Behavioral tests

All the apparatus used for behavioral testing were cleaned with a 70% ethanol solution after each trial to eliminate residual odours. Additionally, for all behavioral tests, the male mice were tested on a different day from the female group. Behavioral experiments were performed from 09:00-17:00.

### Open field test (OFT)

Motion parameters were analyzed using an OFT with an overhead camera (Basler, DE). Behavioral testing was performed in a 50 cm × 50 cm × 50 cm open field arena. Mice were habituated to the testing room for one hour over three consecutive days before testing. On test days, each mouse was placed in the arena and recorded during free movement for 30 minutes. Analysis of total distance, average speed and max speed was performed (ANY-maze software 7.37).

### Walkway

A walkway apparatus developed by the CSMOpto core facility at the University of Calgary was used to automatically record the unsupervised locomotor activity of mice ^76^. Briefly, the apparatus consisted of a clear acrylic arena with an internal walkway accessible through circular holes on two side-entry panels. Motion in the walkway was automatically detected, and video recordings were captured with a monochrome camera (BFS-U3-16S2M-C, FLIR BlackFly S, Richmond, BC, Canada). Mice were habituated to the testing room for one hour over three consecutive days before testing. On test day, each mouse was placed in the arena, and unsupervised recordings of the mouse running through the walkway were automatically recorded. Videos in which mice completed a full walkway run were manually selected for analysis. Gait analysis of the recorded videos was performed using the Visual Gait Lab suite ^36^.The average stride, swing and stance duration were determined for each limb, and average speed was calculated using the stride length and stride duration. Interlimb coordination analysis was performed by using a custom Matlab script for circular statistics.

### Ladder rung

A horizontal ladder rung walking test apparatus was custom-made by the CSMOpto core facility at the University of Calgary, according to Metz and Whishaw ^37,53^. It consisted of two plexiglass side walls (70 cm long and 15 cm high) linked by the insertion of metallic rungs (3 mm in diameter). The ladder was elevated 30 cm above the ground, with a covered home cage at one end. The alley between the walls was set 1 cm wider than the animal’s width to prevent it from turning during the crossing. The task’s difficulty was adjusted by changing the spacing between the ladder rungs. Two different patterns were used: a regularly distributed pattern with rungs spaced at 2 cm (Pattern A) and an irregular pattern with rungs spaced from 1 to 5 cm (Pattern B). Mice were placed on the free end of the ladder rung, and five runs were video recorded using a high-speed camera (Basler, DE). Subsequent analysis was performed using BORIS software 8.20.4 ^77^. The total number of steps and errors were counted. The number of foot faults per limb was scored for each run and averaged.

### In vitro spinal cord preparation

Postnatal mice were anesthetized via hypothermia. Animals were rapidly decapitated and eviscerated to expose the spinal column. The preparation was pinned ventral side up in a silicone elastomer (Sylgard) - coated dissecting dish perfused with carbogenated (95% O_2_-5% CO_2_) artificial cerebrospinal fluid (aCSF) containing (in mM): 4 KCl, 128 NaCl, 1 MgSO_4_, 1.5 CaCl_2_, 0.5 Na_2_HPO_4_, 21 NaHCO_3_, 30 ᴅ-glucose, 310-315 mOsm.), at room temperature (∼21°C). The spinal cord was isolated by performing a ventral laminectomy, and the ventral and dorsal roots (VRs) were sectioned distally. The isolated spinal cord was transected at mid-thoracic segment and was gently removed from the vertebral column. The isolated spinal cord was then transferred to a recording chamber perfused with carbogenated aCSF and placed ventral side up. The bath temperature was gradually increased and maintained at 27°C, which falls within the lower end of the physiological temperature range for neonates ^78^ and helps avoid fluctuations in room temperature. The spinal cord was allowed to recover for one hour before experiments were performed.

### Electrophysiology

Motoneuron extracellular neurograms were recorded from the ventral roots with tight-fitting suction electrodes fashioned from polyethylene tubing (PE50). Generally, neurograms were recorded from the left and right lumbar L2 (L2) and L5 ventral roots. Signals were pre-amplified (10X) and by second stage amplifiers (Cornerstone EX4-400 Quad Differential Amplifier) at 100X for a total amplification of 1000X, with a bandwidth filtering (0.1- 1000 KHz). Amplified signals were acquired in DC and digitized at 2.5 kHz (Digidata 1440A/1550B; Molecular Devices; Sunnyvale, CAs). Data was acquired in Clampe× 10.7 software (Molecular Devices) and saved on a desktop computer for offline analysis.

### Fictive locomotion

Fictive locomotor activity was induced by bath application of N-Methyl-D-aspartic acid (NMDA: 7 µM; Sigma-Aldrich), serotonin hydrochloride (5-HT: 10 µM; Sigma-Aldrich), and dopamine hydrochloride (DA: 50 µM; Sigma-Aldrich) ^25^. Within 5-10 minutes of bath application, these drugs produced alternating neurogram activity. A 10-minute time segment, 30 minutes following drug administration, was extracted for subsequent analysis.

### Electrical stimulation and recording of dorsal root potentials

Dorsal root potentials (DRPs) were recorded with suction electrodes placed at the lumbar L5 dorsal root entry zone. These potentials were evoked by electrical stimulation (0.1 Hz, 0.2 ms square pulses, 1- 20 µA) of either Tibial nerve (Tib) or the distal dorsal root. For each stimulus intensity, six traces were averaged. DRPs parameters, including peak amplitude, AUC, and time to peak, were normalized to the mean value obtained from control animals at a reference stimulus intensity of 1 µA. This was the threshold to elicit a DRP in all animals tested. Normalized values are presented as fractions of the control.

### Data analysis

Locomotor data was processed directly from Clampex files imported into SpinalCore software. The digitized data was filtered (60 HZ notch, 10 Hz low-pass filter) before being rectified. Locomotor rhythm frequency and power were analyzed using auto-wavelet spectral analysis for single ventral root recordings and cross-wavelet analysis for paired ventral root recordings using Spinal Core software ^79^. Phase relationships between rhythms were analyzed using circular statistics and illustrated as circular plots, with phase values normalized to a 0-1 range ^80^. In these plots, the mean vector length (arrow length) reflects how tightly phase values are clustered, indicating consistency of coupling between rhythms, whereas the direction of the arrow represents the mean phase relationship.

### Immunofluorescence Staining

Mice were deeply anesthetized with isoflurane (2%) and transcardially perfused with 0.1M phosphate-buffered saline (PBS), followed by 10% neutral buffered formalin (NBF). The brains were extracted then post-fixed in 10% NBF overnight at 4℃ followed by incubation in 30% sucrose (w/v in PBS) overnight at 4℃ for cryoprotection. 50 µm coronal cryosections were serially collected using a cryostat (Leica Biosystems, ON, CA, Cat. #CM1850 UV). After three washes in 0.1M phosphate buffer (PB), the sections were permeabilized in PB-T blocking solution (0.1% Triton-X 100 in 0.1M PB, Sigma-Aldrich, St. Louis, MO, USA) for 30 minutes, followed by incubation in Blocking PB-T solution (10% donkey serum in 0.1% PB-T) for one hour. Working PB-T buffer (1% donkey serum in 0.1% PB-T) was used in subsequent antibody incubations. The sections were subsequently incubated with a mixture of mouse anti-NeuN (1:500; MAB377, Millipore Sigma) and rabbit anti-FMRP (1:200; 7104S, Cell Signaling) antibodies overnight at 4°C. After two 15-minute washes in working PB-T buffer, the sections were subsequently incubated with a mixture of Alexa 488 donkey anti-mouse (1:1000; A21202, Invitrogen) and Alexa 546 donkey anti-rabbit (1:1000; A10040, Invitrogen) antibodies for 2 hours at room temperature. Controls were included where the primary antibody was omitted to check for non-specific binding of the secondary antibodies. Free-floating brain sections were then mounted onto Superfrost^TM^ slides in VECTASHIELD^®^ PLUS Antifade Mounting Medium (Vector Laboratories Inc., Newark, CA, USA, Cat. #VECTH190010) and cover-slipped. Images were acquired via a confocal laser scanning microscope system (Leica STELLARIS 5). Offline image processing including maximal intensity projections was conducted in Fiji.

### Statistical analysis

Statistical analysis was performed using GraphPad Prism 9.5 (GraphPad Software, Boston, MA). The data was tested for normality. For normally distributed data, an unpaired t-test, one-way, or a two-way repeated-measures analysis of variance (ANOVA) was employed. A mixed model analysis was employed to handle missing values. When assumptions of normality or equal variance were not met, nonparametric tests were conducted. Significant effects detected by ANOVA led to post-hoc analyses (Parametric: Šídák; Non-parametric: Dunn’s), with significance set at *p* < 0.05. For circular statistics, significance was computed using Rayleigh’s test (*p* < 0.05). In some instances, to test for differences in the mean phase across conditions, the Harrison-Kanji test, a circular statistic equivalent of a t-test or ANOVA ^80^, was used. Data were presented as mean standard deviation ± (SD).

## Resource availability

### Lead contact

Further information and requests for resources should be directed to and will be fulfilled by the lead contact, Dr. Patrick J. Whelan.

### Materials availability

There were no new materials generated from this study.

### Data and code availability

Data and Matlab scripts available on request.

## Notes

### Competing Interest Statement

The authors have declared no competing interest.

